# Inferring Multiple Metagenomic Association Networks based on Variation of Environmental Factors

**DOI:** 10.1101/2020.03.04.976423

**Authors:** Yuqing Yang, Xin Wang, Kaikun Xie, Congmin Zhu, Ning Chen, Ting Chen

## Abstract

Identifying significant biological relationships or patterns is central to many metagenomic studies. Methods that estimate association networks have been proposed for this purpose, but they assume that associations are static, neglecting the fact that relationships in a microbial ecosystem may vary with changes in environmental factors, which can result in inaccurate estimations. We propose a computational model, k-Lognormal-Dirichlet-Multinomial model (kLDM), which estimates multiple association networks that correspond to specific environmental conditions according to values of environmental factors (EFs), and simultaneously infers microbe-microbe and environmental factor-microbe associations for each network. We showed the effectiveness of kLDM on synthetic data, a colorectal cancer dataset, as well as the TARA Oceans and American Gut project datasets. The results showed that the widely used Spearman’s rank correlation coefficient (SCC) performed much worse than other methods, indicating the importance of separating samples by environmental conditions. We compared cancer fecal samples with cancer-free samples, and our estimation showed fewer associations among microbes but stronger associations between specific bacteria such as five colorectal cancer (CRC)-associated OTUs, indicating gut microbe translocation in cancer patients. Some environmental-factor-dependent associations were found within marine eukaryotic community, and gut microbial heterogeneity of irritable bowel disease (IBD) patients was detected. Results demonstrated that kLDM could successfully unravel the underlying biological associations. In summary, our study presents a computational framework that can elucidate the complex associations within microbial ecosystems. The kLDM program, R, and python scripts, together with all experimental datasets are all accessible at Github (https://github.com/tinglab/kLDM.git).

## Introduction

Microbes interact closely with the human body and the environment human lives, which affect our health and ecosystem by changing their microbiome and metabolome [1–3]. Metagenomic high-throughput sequencing technology plays a vital role in the study of microbes inhabiting various human body sites and different natural ecological environments. Such tool allows us to analyze microbial constituents of the microbiota and microbiome and interactions within a particular microbial community among microbes and between microbes and environmental factors (EF). The environmental factors, such as states of human diseases, genotypes of hosts, values of some physiological and biochemical indicators, quantization of people’s lifestyle factors, and concentrations of nutrients, are known to be associated with variation of abundance of microbes [4–8]. Recently, we have observed rapid increase of large scale metagenomic studies aiming to discover biological interactions [9,10], specifically, how microbes interact with other microbes and EFs.

Biological interactions can be considered as positive and negative relationships among microbes and between EFs and microbes, which can be inferred indirectly by predicting patterns of variation of microbial counts and values of EFs using association inference methods. A positive relationship may imply commensalism among microbes or microbial dependency on some EFs, whereas a negative relationship can be parasitism, competition among microbes, or inhibition of EFs on microbes [11]. Furthermore, interactions among microbes and between EFs and microbes depend on ***environmental conditions* (EF conditions)**, which, in this article, are defined as some specific ranges of values of EFs. Under similar EF conditions, interactions in the microbial community can be stable, while they can change dynamically with the alteration of EF conditions (**Figure 1A**). For example, marine microbial interactions may vary by season and depth [12], and interactions among gut microbes can change according to the onset of diseases and disease states [4,13]. In these two examples, the environmental conditions are specific seasons and ocean depths, and specific diseases and disease states, respectively. Overall, biological interactions are dynamic, a phenomenon that should be considered in any approach to association inference.

**Figure 1.**
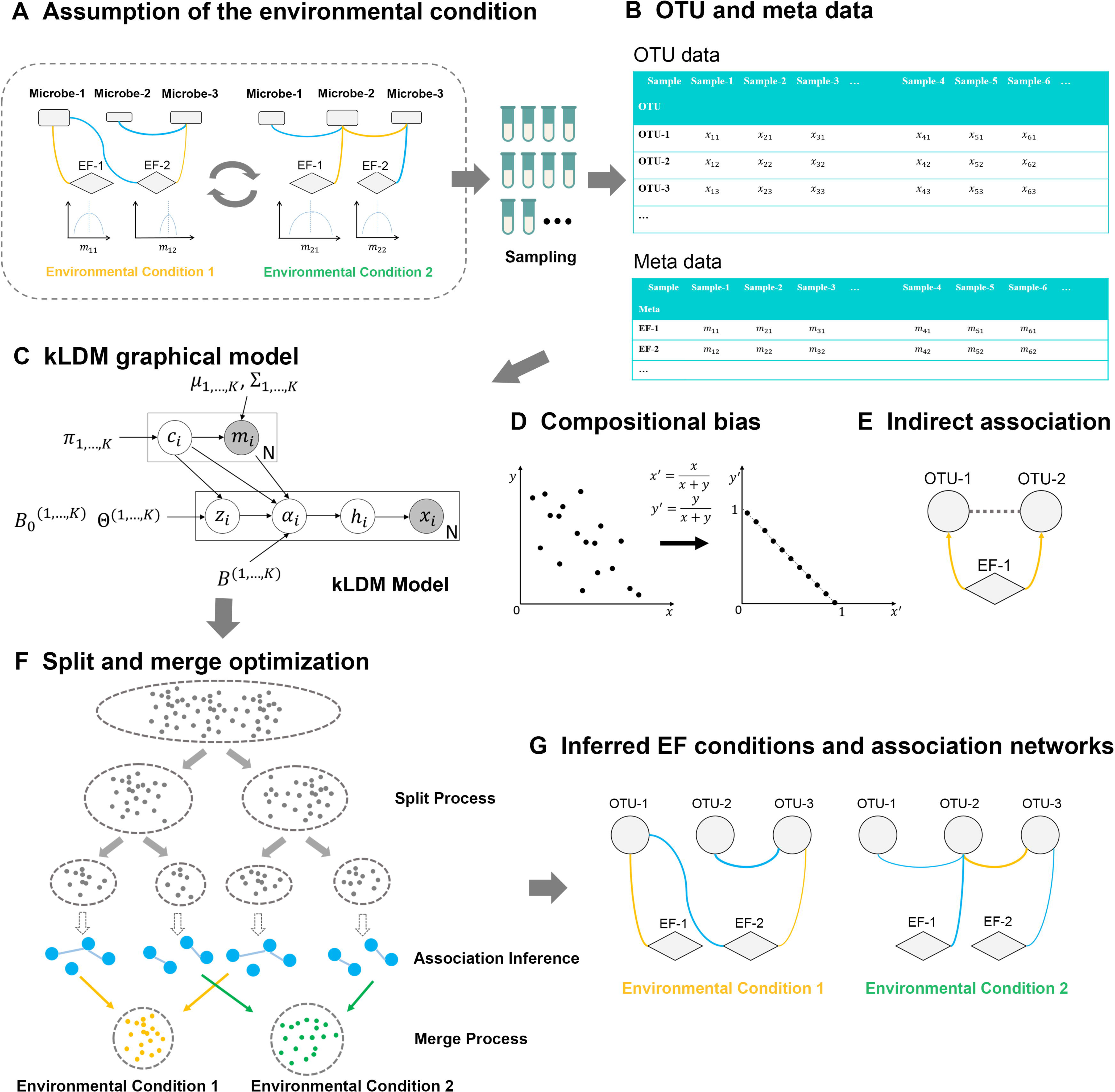
Schema of the kLDM model. **A.** Multiple environmental conditions are assumed to exist in real environment, and the EF condition can change with time. An EF-condition refers to (1) a group of samples where values of EFs fall into a small and defined range, and (2) under this EF condition, interactions within the microbial community are stable. **B.** Sequencing samples with related meta data, possibly from multiple environmental conditions (EF conditions), are collected. After data preprocessing, clustering, and annotating, OTUs’ counts are obtained for each sample. The information about which two samples belong to the same EF-condition is unknown beforehand. **C.** The kLDM graphical model assumes K EF conditions within N samples, and infers the number of EF-conditions and associations among OTUs and associations between EFs and OTUs under every EF-condition. Two matrixes, *B*^*k*^ and Θ^*k*^, record direct EF-OTU associations and OTU-OTU associations on the *k*^*th*^ EF condition, respectively. Vectors *x*_*i*_ and *m*_*i*_ are OTU’s counts and values of EFs in the *i*^*th*^ sample respectively. At every EF-condition, we assume that values of EFs follow a multivariate Gaussian distribution that is, parameterized by *μ*_*k*_ and ∑_*k*_. The rest of the variables include *h*_*i*_, which represents the latent relative ratios of microbes in *i*^*th*^ sample; *α*_*i*_, which is the absolute abundance of microbes; 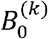, which represents the impact of unknown factors affecting OTUs’ abundance; *c*_*i*_, indicating that the *i*^*th*^ sample belongs to EF condition *c*_*i*_; and *π*_*k*_ which is the mixture weight of the *k*^*th*^ EF condition. **D.** Compositional bias is caused by the normalization process on OTUs’ counts. After normalization, microbial relative abundance sums to one. **E.** The indirect association between OTU-1 and OTU-2 induced by the common environmental factor EF-1 can be recognized by kLDM, which takes the environmental factor-microbe (EF-OTU) association into account. **F.** For kLDM, the number of EF conditions and association networks of every EF condition are estimated by a spilt and merge optimization algorithm. Both values of EFs and associations of microbes are taken into account to determine the EF condition. **G.** Parameters estimated by kLDM can be visualized into environmental conditions and association networks. The blue and yellow edges correspond to negative and positive associations, respectively, and the thickness of an edge is proportional to the association values.

However, environmental conditions are sometimes neither obvious, nor are they known in advance in some studies. For example, the American Gut project[14] collected information on dozens of human lifestyle factors related to individuals’ diet and living habits, as well as metagenomic sequencing data for thousands of individuals. However, it is unclear which subset of these individuals belong to the same environmental condition, which, by definition, not only have similar values of some, or all, of these lifestyle factors but also share identical microbial interactions. Besides, even if we have obtained hosts’ disease states, grouping samples into the cases and controls while ignoring other environmental factors may be rough, because the underlying patterns of microbial compositions and associations may be very complex, consist of many subgroups, and can be aligned with neither the cases nor the controls. These problems call for novel computational methods to discover potential environmental conditions and infer association networks together.

Due to a limited number of sequencing samples in studies, previous association inference methods estimate static association network using all samples [15–20]. These methods can be classified into two categories: those that compute pairwise associations independently, and those that estimate multiple associations simultaneously. The widely used Pearson correlation coefficient (PCC) and Spearman’s rank correlation coefficient (SCC) belong to the former, as well as the local similarity association (LSA) [19]. In contrast, CCREPE[16], SparCC[15], SPIEC-EASI[20], CCLasso[18] and mLDM[17] belong to the latter. The latter methods consider compositional bias [21] caused by the normalization process of microbial read counts, which divides microbial read counts over total sum of read counts to obtain relative abundance. Denote the relative abundance of the *j*^*th*^ OTU as 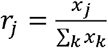 where the *x*_*j*_ is its read count and we get ∑_*k*≠*j*_*Cov*(*r*_*k*_, *r*_*j*_) = −*Var*(*r*_*j*_). It can be seen that the commonly proposed normalization process [22] introduces dependency into microbial relative abundance, and as such, it reduces the efficiency of association inference (Figure 1D). CCREPE, SparCC and CCLasso estimate OTU-OTU (microbe-microbe) correlations by computing covariance among microbes, while SPIEC-EASI and mLDM infer conditionally dependent OTU-OTU associations by estimating the precision matrix among microbes. With the exception of mLDM, none of these methods considers associations between EFs and microbes. Taking both compositional bias and the large variance of read counts into account, our previous work, mLDM [17] estimates both OTU-OTU and EF-OTU associations more accurately by removing indirect associations among microbes induced by common EFs (Figure 1E).

However, all methods noted above assume only one biological network, neglecting that biological interactions can be different with the change of EFs. In this case, if samples from two different environmental conditions are combined to infer associations, the results will reflect the change of environmental conditions, instead of real associations under these two environmental conditions, which may lead to false associations and conclusions. For example, association networks of gut microbes in patients with liver cirrhosis and healthy individuals are distinctly different [23], and to predict gut microbial interactions of disease patients, samples from the healthy population should be excluded.

In order to estimate OTU-OTU associations and EF-OTU associations conditional on environmental conditions, we propose a new hierarchical Bayesian model, k-Lognormal-Dirichlet-Multinomial model, called kLDM (Figure 1), which can determine the number of environmental conditions automatically and infer complex associations simultaneously. Under every EF condition, kLDM considers the compositional bias and large variance of read counts, and it estimates both conditionally dependent OTU-OTU associations and direct EF-OTU associations. Most significantly, associations that vary according to environmental conditions can be elaborated by kLDM. It should be noted that the direct or conditionally dependent associations are mathematical concepts that approximate biological interactions, rather than indicating direct biological interactions or causal relationships. In addition, considering sequencing of marker genes such as 16S rRNA/18S rRNA genes is widely used to investigate microbial compositions of samples, kLDM is designed for data produced by such marker gene sequencing technology. To our best knowledge, kLDM is the first method to infer multiple association networks based on variation in the values of environmental factors. Here, we validate the efficiency and robustness for sparse and compositional data of kLDM on synthetic datasets via comparisons with state-of-art approaches, then apply it to colorectal cancer (CRC) data to show its ability to find well-defined EF conditions, and explore its application on TARA Oceans data and American Gut project data to discover potential EF conditions and novel biological relationships.

## Materials and methods

### Materials used for kLDM evaluation

#### Gut microbial samples from Colorectal Cancer study

OTU table and meta data of metagenomic gut 16S rRNA sequencing data of healthy individuals and patients with Colorectal Cancer (CRC) were obtained from Baxter et al. [13]. We constructed a dataset which consisted of 5 known CRC-associated OTUs (*Peptostreptococcus*, *Parvimonas*, *Fusobacterium*, *Porphyromonas*, and *Prevotella*) reported in the original article, 112 common OTUs, as observed in more than half of all 490 samples, and 4 EFs, namely FIT result, age and gender, and diagnostic states of donors. The ‘FIT’ result refers to the positive or negative result of fecal immunochemical test. The recorded diagnostic states include ‘Normal’, ‘High Risk Normal’, ‘Adenoma’, ‘Advanced Adenoma’, and ‘Cancer’, all of which are determined by colonoscopy examination and review of biopsies, were also recorded. The diagnostic states were modeled as five numerical values from 1 to 5, with a higher value representing a more serious disease state.

#### TARA Oceans eukaryotic dataset

In the study of **TARA Oceans** project about marine eukaryotes [24], read counts of OTUs were obtained by sequencing the V9 region of 18S rRNA genes of eukaryotes, and 91 genus-level matched eukaryotic symbiotic interactions were summarized. Because of the limited number of samples, after filtering OTUs that exist in < 40% of samples and getting rid of samples with missing EFs or abnormal counts, a subset including 67 OTUs related to genus-level symbiotic interactions and 17 EFs from 221 samples was constructed. The 17 EFs are the depth of water, chlorophyll maximum, maximum of Brünt-Väisälä frequency, maximum dissolved oxygen, and minimum dissolved oxygen; concentrations of salinity, oxygen, phosphate, silica, and chlorophyll; temperature, sunshine duration, moon phase, maximum Lyapunov exponent, residence time, latitude and longitude.

#### 16S rRNA sequencing samples from American Gut project

OTU table and meta data from the American Gut project[14] were downloaded from the ftp site (ftp://ftp.microbio.me/AumericanGut), and 22 meta data about individuals’ diet and living habits were selected. Among 22 meta data, three factors were associated with living habits (alcohol, exercise, and smoking frequencies), the remaining 19 factors were connected with diet (frequencies of fermented plant, frozen dessert, fruit, high fat red meat, home-cooked meals, meat eggs, milk cheese, milk substitute, olive oil, probiotic, red meat, salted snacks, seafood, vegetable, vitamin D supplement, vitamin B supplement, whole grain, and whole eggs). The value of meta data can be categorized as one of ‘Never’, ‘Rarely (less than once/week)’, ‘Occasionally (1-2 times/week)’, ‘Regularly (3-5 times/week)’ and ‘Daily’. For convenience, we recode these categories into integers from 1 to 5 according to their frequencies. OTUs are filtered based on their average size and non-zero times in all samples. Samples with huge or small numbers of reads, as well as those with evenness < 2, are removed. Finally, 11,946 samples with 216 OTUs and 22 EFs are obtained. The python script to process the OTU .biom file can be found at the Github (https://github.com/tinglab/kLDM.git). Information on individuals’ disease status was also extracted, and samples with diseases were labeled, including cardiovascular disease, small intestinal bacterial overgrowth, mental illness, lactose intolerance, diabetes, inflammatory bowel disease (IBD), irritable bowel syndrome, *C. difficile* infection, cancer, and obesity.

### Assumptions of kLDM hierarchical Bayesian model

The kLDM model assumes that interactions among microbes are regulated by environmental factors and that they tend to be constant when environmental conditions are identical, but may change as environmental conditions change (Figure 1A). Or in other words, kLDM accounts for variation in the values of EFs. In the same environmental condition, EF values fluctuate, but only within a small range, and core microbes stay the same, their associations remaining stable. However, when environmental conditions vary, values of EFs, microbial species, and their associations can all change. In addition, considering that distribution of environmental conditions may be continuous and complex [25], kLDM uses mixtures of multiple clusters with known distributions to approximate and capture main patterns, and each cluster can represent an environmental condition.

### The hierarchical structure of kLDM model

Let 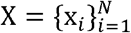 be N sequencing samples where *x*_*i*_ ∈ ℕ^*P*^ is the *i*^*th*^ sequencing sample with P microbes, and *x*_*ij*_ corresponds to the read count of the *j*^*th*^ microbe or Operational Taxonomic Unit (OTU). Values of environmental factors are represented as 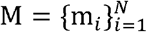, where *m*_*i*_ ∈ ℝ^*Q*^ is a Q-dimensional vector and *m*_*ij*_ records the value of environmental factor j of the *i*^*th*^ sample. kLDM models the EF vector *m*_*i*_ as a multivariate Gaussian distribution, and assumes that EFs and whole data consist of K mixtures of Gaussian components. Thus, samples in which EF vectors belong to the same Gaussian component can be viewed as drawn from the same environmental condition and share identical OTU-OTU and EF-OTU association networks. K environmental conditions are denoted as 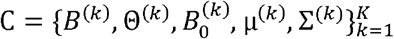, where matrices *B*^(*k*)^ and Θ^(*k*)^ are the associations between EFs and OTUs and among OTUs for the *k*^*th*^ environmental condition, respectively, 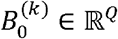 is a basis vector, and μ^(*k*)^ and Σ^(*k*)^ are the mean vector and covariance matrix of EFs for the *k*^*th*^ environmental condition. In addition, weights of K environmental conditions are 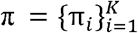, where π_*k*_ is the weight of the *k*^*th*^ EF condition, and 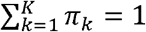.

Figure 1C explains the structure of the kLDM model. x_*i*_ and m_*i*_ represent the vectors of read counts and values of EFs of the *i*^*th*^ sample, respectively. The vector h_*i*_ is a P-dimensional latent variable of which the value h_*ij*_ is the latent relative abundance of the *j*^*th*^ microbe in sample *i* and α_*i*_ can be regarded as the absolute abundance vector corresponding to h_*i*_. kLDM assumes that the absolute abundance α_*i*_ determines the relative abundance h_*i*_ of P microbes during the sample preparation process, and that the microbial read counts vector x_*i*_ is generated based on relative microbial abundance during the sequencing process. Sample *i* can be seen as being collected from a specific environmental condition denoted by *c*_*i*_ with mixture weight 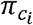. The absolute abundance α_*i*_ is decided by two sources of associations: 1) associations between environmental factors and microbes under the 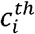 environmental condition, which can be parameterized by a linear term 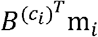; and 2) associations among microbes, the effects of which are included in the EF condition-specific latent variable 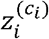. The vector 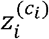 follows a multivariate Gaussian distribution, the parameters of which are determined by the basis vector 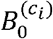 and the precision matrix 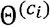, which records OTU-OTU associations at the EF condition *c*_*i*_.

The generative process of the hierarchical Bayesian model, kLDM, is shown,

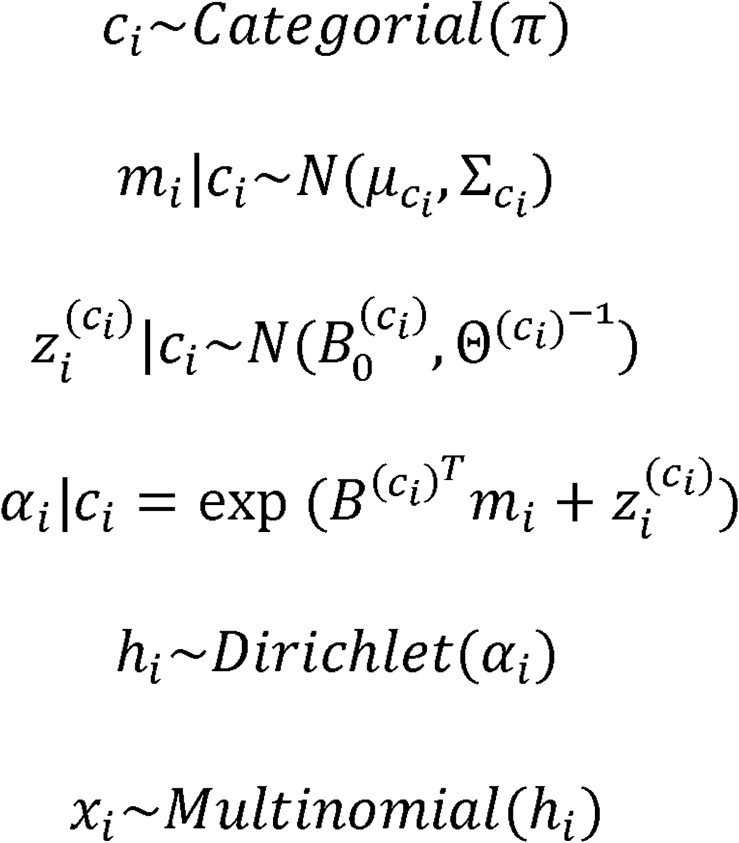

where 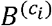 is a Q × P matrix where 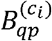 is the association between the *q*^*th*^ environmental factor and the *p*^*th*^ microbe, and 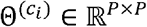 is a P × P inverse covariance matrix. During the sequencing process, millions of DNA molecules are selected randomly from the DNA library; therefore, we let the sequencing counts *x*_*i*_ be a multinomial distribution,

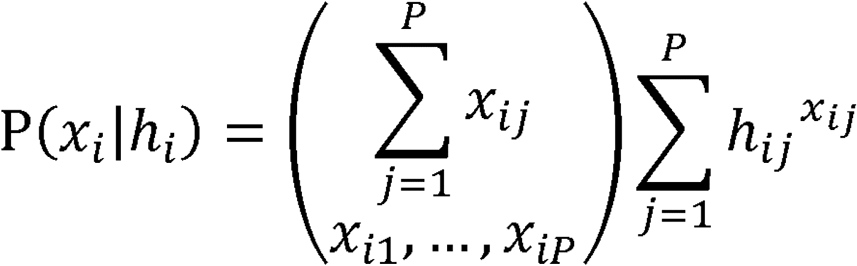

where the relative abundance *h*_*i*_ represents the microbial relative ratios within the DNA library and 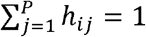. Considering the conjugacy of Dirichlet and multinomial distribution, *h*_*i*_ follows the Dirichlet distribution,

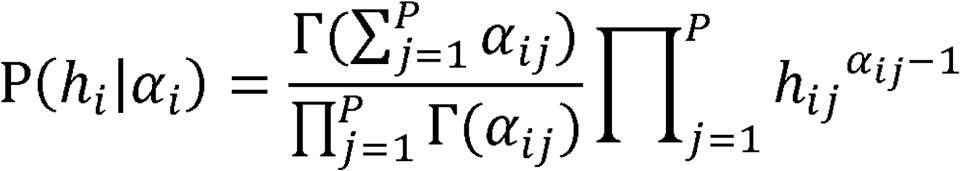

where α_*i*_ is the vector for absolute microbial abundance in the *i*^*th*^ sample, which, in turn, determines the relative ratios in the library. In Dirichlet-multinomial distribution, the covariance of two OTUs’ counts *x*_*ij*_ and *x*_*ik*_ in *i*^*th*^ sanple is 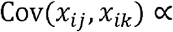 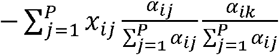, which corresponds to a negative bias, and is regulated by both the sequencing depth 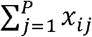 and OTUs’ relative abundance 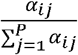. The Dirichlet-multinomial distribution is integrated into the hierarchical structure of kLDM to model the compositional characteristics of sequencing count data.

Assuming that microbes in the *i*^*th*^ sample come from environmental condition c_*i*_, their absolute abundance α_*i*_ is affected by both environmental factors m_*i*_ and interactions within the community. We model this into a lognormal distribution which is suitable for most microbial abundance [26,27]:

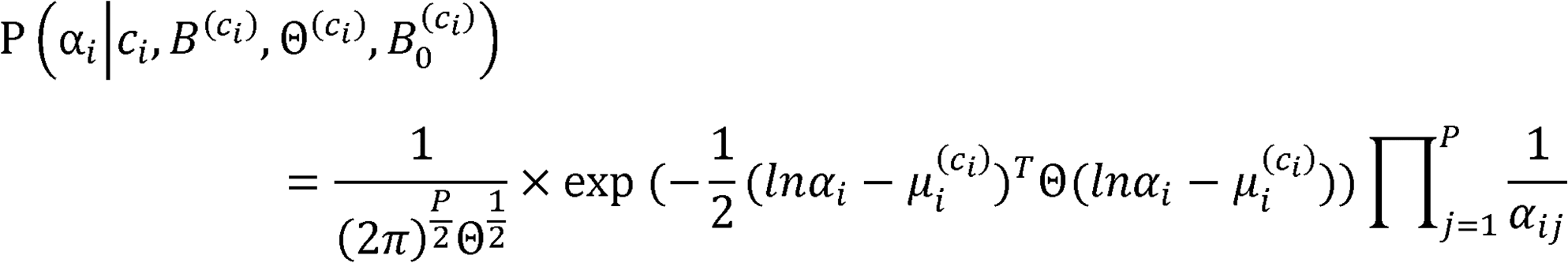

where 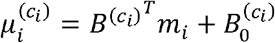. This can be simplified into the following formula using the relationship between the lognormal and normal distributions,

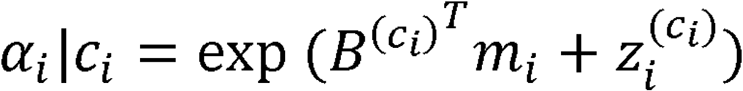

where 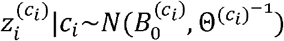.

Under the 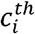 environmental condition, *m*_*i*_ follows the multivariate Gaussian 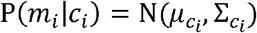 and the weight of the 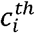 EF condition is set to the categorical distribution with the parameter π as,

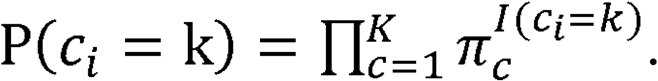

We are interested in the parameters *B*^(*k*)^, Θ^(*k*)^(*k* = 1,…, *K*), where 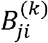 infers direct association between the *i*^*th*^ microbe and *j*^*th*^ environmental factor at the *k*^*th*^ environmental condition, and 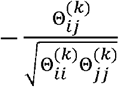 is the conditionally dependent association between the *i*^*th*^ and *j*^*th*^ microbes (Figure 1G).

### Sparse association inference of kLDM in theory

The generative model can be solved theoretically via Expectation-Maximization (EM) algorithm and maximum a posteriori (MAP) estimation for the latent variable 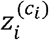. We assume that associations among microbes and between environmental factors and microbes are sparse and can be inferred by kLDM with sparsity constraints. Detailed formulas of inference and optimization can be found in the **Supplementary Information**.

However, two potential problems confront this theoretical sparse association inference, thereby limiting the practicality of the model. First, results are very sensitive to the initialization of parameters because the EM algorithm can converge to a local minimum. Second, estimating the number of EF conditions is very time-consuming. Therefore we explored more effective approaches and finally adopted an efficient split-merge algorithm as detailed below.

### Implementation of split and merge Algorithm for kLDM

kLDM adopts a split-merge algorithm to estimate the number of environmental conditions and sparse OTU-OTU and EF-OTU associations under each environmental condition [28]. First, we partition samples into fine-grained clusters using values of environmental factors, such that samples within a cluster can be regarded as belonging to the same environmental condition. Second, since this partition is not perfect, we merge multiple clusters into one if they share similar environmental conditions and association networks. The final output is a set of sample groups, each with distinct predicted association networks.

More specifically, the split process starts from a single cluster with all samples, and then iteratively selects a cluster and partitions it into two clusters until the number of samples in each cluster is smaller than a pre-defined threshold *N*_*min*_. This process results in the construction of a binary tree for environmental factors, each node of which corresponds to one cluster of the samples, and each leaf node of which corresponds to a set of samples with similar values of environmental factors. We assume that EF vector *m*_*i*_ follows the multivariate Gaussian distribution, and when a cluster is split in two, two Gaussian mixtures are used to model the distribution of EFs of the cluster as,

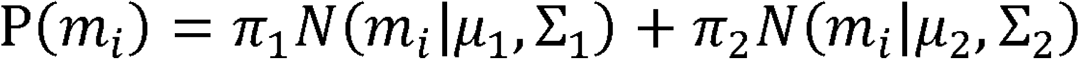

where *π*_1_ + *π*_2_ = 1 and *π*_*j*_(*j* = 1,2) is the weight of the *j*^*th*^ component. After estimating these parameters via the EM algorithm, we re-assign each sample of the original cluster to one of the two new clusters with the larger posterior probability.

Then association networks are estimated for the leaves via mLDM [17], which is similar to kLDM when the number of the environmental condition is set to one (K = 1). We re-implemented the mLDM algorithm with C++ and OpenMP to improve its stability and efficiency, and a comparison of running time and memory usage by kLDM and mLDM on single association network inference was shown in Table S1. Inferred associations will be used for the merge process.

The merge process aims to recover clusters being partitioned into multiple leaf nodes caused by the greedy approach of the split process. The merge process adopts a bottom-to-top strategy, starting from internal nodes at the lowest level and traversing up to the root, to identify leaf nodes for merging. For each internal node, the algorithm traverses down to its left and right branches to search for leaf nodes. The two leaf nodes with the smallest inter-cluster distance, as measured by the Euclidean distance between the mean values of EFs, are merged. kLDM will estimate the associations and the EBIC score for the merged cluster. If its EBIC score is smaller than the sum of the EBIC scores of the two leaves, then the merged cluster is kept and substitutes the one that has closer mean values of EFs, while the other is discarded. Otherwise, the merged cluster is discarded. This split-merge process reduces the time complexity by limiting the operation at each step to partition one large cluster into two, or merge two nearby clusters into one. The algorithm can run in parallel, making kLDM very efficient in practice.

### Synthetic data generation process

We generate synthetic datasets by specifying the number of microbes, environmental factors, and clusters, as well as the range of the number of samples per cluster. We then construct samples for every cluster via a generative process of the kLDM model separately. For the *i*^*th*^ cluster (i = 1,2,…,K), the EF-OTU association matrix *B*_*i*_ is produced by sampling from the interval [−0.5,0.5] uniformly, with only 15 percent of the elements set to non-zero. The OTU-OTU association matrix Θ_*i*_ is generated using the R package ‘huge’ [29], which outputs a precision matrix for which the adjacency matrix can be random, cluster, scale-free, hub or band graph. Every graph corresponds to a specific association structure among microbes. Values of the mean vector of the environmental factor of the cluster can be obtained by sampling uniformly from the interval [i, (i + 0.5) ∗ i]. Then the Dirichlet-multinomial samplers are produced using the R package ‘HMP’. For parameters of every setting, ten repetitions are conducted to generate synthetic data and then the mean and standard deviation of evaluation metrics are calculated for comparison. Public R language codes of CCLasso and SPIEC-EASI are used. The implementation of SparCC is provided by SPIEC-EASI. SCC and SCC(all) are implemented directly in built-in functions of R language. The ‘mb’ (meinshausen-buhlmann) estimation method is set for SPIEC-EASI, and default parameters of CCLasso, SPIEC-EASI, and SparCC are utilized. In addition, the p-value is set to 0.05 for SCC and PCC to save significant associations.

### Evaluation metrics on synthetic data

ROC curves and AUC scores are utilized to evaluate the performance of association inference. Every cluster estimated by kLDM is shown by two ROC curves, the ROC curve of the OTU-OTU associations and that of the EF-OTU associations. AUC scores are the area under ROC curves. When we compute AUC scores, signs of estimated associations are neglected. As for thresholds to compute AUCs, the absolute values of calculated associations are selected for kLDM, SparCC, CCLasso, and SPIEC-EASI, and p-values are used by SCC and PCC. When plotting ROC curves for SCC and CCLasso, calculated coefficients are compared with the true correlation matrix, as deduced by the inverse of the precision matrix Θ_*i*_. One estimated nonzero association is regarded as a true positive if its value is also nonzero in ground truth. If values of one association are both zero in inferred and real results, the association is a true negative.

## Results

### Comparing kLDM with other methods on synthetic datasets

In the following, we first assess the performance of kLDM using synthetic datasets, and then we evaluate how similarity of underlying networks and missing information, i.e., EFs, affect the performance of kLDM.

Detailed experiments are conducted to show the effectiveness of kLDM by comparing it with Spearman’s Correlation Coefficient (SCC), CCLasso, and SPIEC-EASI. These three models are included because they were tested in our previous studies [17] and showed advantages over other methods. SCC estimates both EF-OTU and OTU-OTU associations, and CCLasso solves the covariance matrix among microbes, while SPIEC-EASI performs well in inferring the precision matrix among OTUs. For fair comparison, since none of them considers more than one network, we partition data into clusters according to the results reported by kLDM and apply these programs on each cluster to infer associations. It should be noted that the results of SCC using all samples, denoted as SCC(all), are plotted as the baselines. The ROC curve and AUC score on every cluster are used to compare the performance.

First, we examine the relationship between the number of samples per cluster N, and the efficiency of kLDM by setting the following parameters: K = 2 clusters, P = 50 microbes, Q = 5 environmental factors, and N samples with two ranges denoted as [*N*_*min*_, *N*_*max*_] = [100,200] and [200,400]. These two settings of N have identical associations of microbes and between environmental factors and microbes. **Figure 2** shows that the ROC curves and AUC scores of kLDM are consistently better than those of the other methods, and the power of kLDM on recovering OTU-OTU and EF-OTU associations increases with increased N. However, we observe that the ROC curves of SCC (all) in most situations are lower, showing the importance of separating samples by environmental conditions. It should be noted that SPIEC-EASI does not estimate association efficiently due to its strict model selection. Samples from different EF conditions may introduce many noises and disturb results of SPIEC-EASI. Besides, sensitivities and specificities of top associations estimated by the five methods were compared using the synthetic dataset with K=2, P=50,Q=5 and N∈[100,200] corresponding to Figure 2A. From Table S2, we observe that kLDM infers OTU-OTU or EF-OTU associations with higher sensitivity and superior specificity.

**Figure 2.**
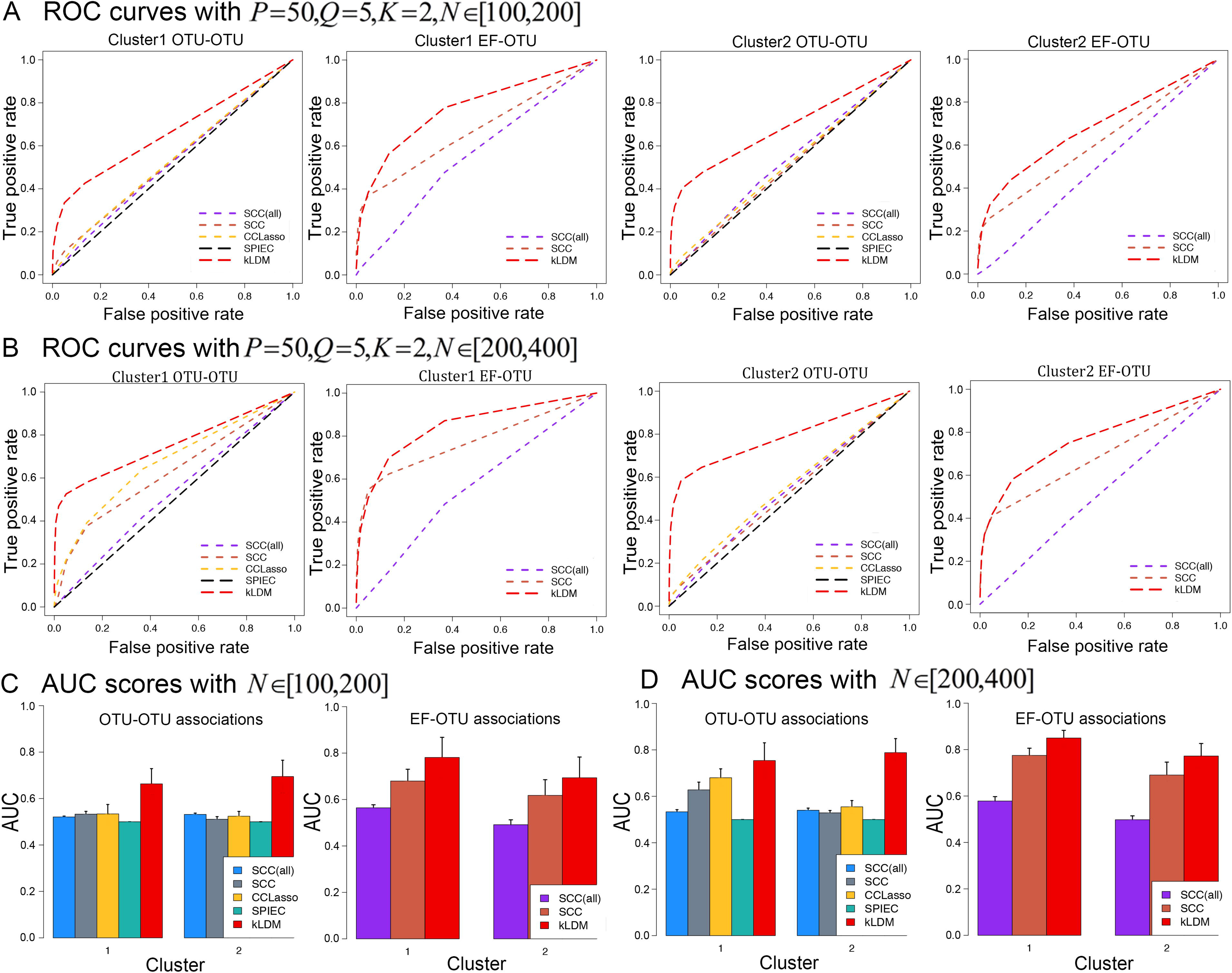
Comparing ROC curves and AUC scores of kLDM with other methods on synthetic data. ‘P’, ‘Q’ and ‘N’ represent the numbers of OTUs, EFs and samples per cluster, respectively. **A.** Receiver operating characteristic (ROC) curves with 50 OTUs, 5 EFs and 2 clusters. The number of samples per cluster is set to [100, 200]. Two clusters’ ROC curves on OTU-OTU and EF-OTU associations are plotted orderly. The red line corresponds to results of kLDM. **B.** ROC curves with the same parameters as **A** except that the size of clusters is set to [200, 400]. **C.** Area under the curve (AUC) scores for **A**. Scores of OTU-OTU associations and EF-OTU associations for two clusters are shown. **D.** AUC scores for **B**.

Then, the scalability of kLDM was investigated with P=100 and 200 microbes. The results are plotted in Figure S1A and S1B. In both situations, the AUC scores of kLDM are all higher than those of the other approaches, which verifies its fine scalability, owing to the re-implementation of mLDM with C++ language and utilizing parallel computing. Next, we test kLDM by increasing the number of clusters to 3 and 4 (Figure S1C and S1D), and again kLDM achieves the best results.

Because the split process of kLDM is affected by similarity of environmental factors between clusters, we only change the distance between mean values of environmental factors of two clusters and keep other parameters unaltered to examine the performance of kLDM. From Table S3, as expected, when the distance between two EF mean vectors is large enough, such as 1.5 or 2.0 denoted as ‘***1.5 baseline***’ and ‘***2.0 baseline***’, kLDM infers associations accurately. However, the effectiveness of kLDM reduces when the distance becomes smaller, such as 1.0 (‘***1.0 baseline***’), especially on the inference of OTU-OTU associations. This can be attributed to the tendency of kLDM to group samples together to infer common associations when two clusters have similar values of environmental factors, but different associations.

To test the effect of similarity of EF-OTU or OTU-OTU associations among clusters on the performance of kLDM, we generate two new datasets for each distance (‘1.0, 1.5 and 2.0 baseline’) by setting the *B*_*i*_ or Θ_*i*_ of two clusters (i=1,2) equal separately (‘same EF-OTU’ or ‘same OTU-OTU’ in Table S3). Compared with results of the corresponding ‘baseline’, AUC scores of ‘same EF-OTU’ and ‘same OTU-OTU’ do not change much. From this, we concluded that the similarity of EFs influences kLDM more than the similarity of associations among environmental conditions.

In simulated experiments, all EFs are used to estimate association networks, which may not be feasible in reality because some EFs can be missing. Therefore, we simulate new datasets with only partial EFs to test kLDM’s effectiveness. In Figure S2, AUC scores of kLDM on two clusters are shown with different proportions (20%, 40%, 60% and 80%) of EFs retained to infer association networks. With subsets of EFs, the AUC scores of kLDM decline. This indicates the importance of including as many EFs as possible. Furthermore, with only 60% EFs, kLDM achieves results comparable to those of CCLasso and SCC with the whole dataset, which again shows the superiority of kLDM (Table S4).

### kLDM captures variation in associations of gut microbiota of colorectal cancer patients

The relationship between human gut microbiota and colorectal cancer has been explored in previous studies [30–32]. We choose the metagenomic 16S rRNA sequencing dataset from Baxter et al. [13], consisting of 117 OTUs and 3EFs, and validate the efficiency of kLDM on capturing the variation of associations in the microbial community. kLDM reported two clusters: Cluster 1 annotated as ‘Cancer’, which includes 90% of ‘Cancer’ samples with significantly higher positive FIT values (P-value 7.18e-22) (Table S5), and Cluster 2 denoted as ‘Healthy’, which contains 83.7% of healthy samples (‘Normal’ and ‘High Risk Normal’) (Table S6). ‘Adenoma’ and ‘Advanced Adenoma’ samples exist in both Cluster 1 and Cluster 2. The clustering result of kLDM does not split samples simply according to the diagnostic state; instead, it takes all EF values, abundances and associations of microbes into account.

Different patterns of microbial abundance and associations are found between these two clusters. *Prevotella* is abundant in samples from both clusters, while the other four CRC-associated OTUs are significantly over-represented (P-value < 1e-3) in samples of Cluster 1 (Figure S3). Comparing OTU-OTU association networks between Cluster1 and Cluster2, as shown in Figure S4, most gut microbes of ‘Healthy’ patients (Figure S4B) connect quite closely, and the distribution of associations among microbes is balanced, while few associations are observed between the aforementioned 5 known CRC-associated OTUs and other microbes. In contrast, associations among gut microbes of ‘Cancer’ patients’ (Figure S4A) are relatively sparse, and strong correlations are found between specific bacteria (**Table 1**). *Peptostreptococcus, Parvimonas, Fusobacterium*, and *Porphyromonas* show strong correlations with each other, but don’t connect with other microbes, while *Prevotella* is more associated with other common OTUs, such as *Phascolarctobacterium* and *Clostridium_XlVa*. Based on the difference of microbial abundance and associations’ distribution within two clusters, the gut microbiota translocation in cancer samples can be found, and *Prevotella* may play a specific role in tumorigenesis.

**Table 1.** Top associations of CRC-associated microbes for two clusters of CRC dataset found by kLDM.

| Cluster 1 (Cancer) |  |  |
| --- | --- | --- |
| OTU | EF or OTU | Association |
| *Parvimonas* | *Porphyromonas* | +0.238 |
| *Prevotella* | *Phascolarctobacterium* | -0.234 |
| *Peptostreptococcus* | *Porphyromonas* | +0.220 |
| *Peptostreptococcus* | *Parvimonas* | +0.191 |
| *Parvimonas* | *Fusobacterium* | +0.180 |
| *Porphyromonas* | *Prevotella* | +0.173 |
| *Peptostreptococcus* | *Fusobacterium* | +0.155 |
| *Fusobacterium* | *Porphyromonas* | +0.155 |
| *Porphyromonas* | *Akkermansia* | +0.121 |
| *Prevotella* | *Clostridium_XIVa* | -0.095 |
| *Prevotella* | *Clostridium_sensu_stricto* | +0.091 |
| *Prevotella* | *Clostridium_IV* | -0.090 |
| *Parvimonas* | *Prevotella* | +0.085 |
| *Prevotella* | *Clostridium_XIVa* | -0.082 |
| *Porphyromonas* | <u>Diagnostic State</u> (EF) | +0.294 |
| *Fusobacterium* | <u>FIT</u> (EF) | +0.245 |
| *Peptostreptococcus* | <u>Diagnostic State</u> (EF) | +0.281 |
| **Cluster 2 (Healthy)** |  |  |
| **OTU** | **EF or OTU** | **Association** |
| *Prevotella* | <u>Diagnostic State</u> (EF) | +0.274 |

*Note: 'CRC' is the abbreviation for colorectal cancer. Results of CRC-related* *OTU-OTU and EF-OTU associations, selected from top 1% weighted associations* *from the two clusters. Five microbes related to CRC, as reported in previous study,* *are labeled in bold and the environmental factors are underlined. The 'Diagnostic* *State' consists of 'normal', 'high-risk normal', 'advanced adenoma', 'adenoma' and* *'cancer' states. 'FIT' stands for fecal immunochemical test results.*

Previous studies have shown that *Peptostreptococcus* and *Fusobacterium* are associated with inflammation [33,34]. Our results also confirm that *Peptostreptococcus* is positively correlated with *Fusobacterium* in ‘Cancer’ patients and that it is over-represented in CRC fecal samples. From Table 1, *Porphyromonas* and *Peptostreptococcus* show positive correlations with diagnostic states in Cluster 1, suggesting the significance of these two bacteria in CRC diagnosis. On the other hand, *Prevotella* is positively associated with the diagnostic states in Cluster 2, indicating a high predictive value, which could be a useful indicator for CRC diagnosis. Research has shown that *Fusobacterium* may act as a passenger microbe to perpetuate tumorigenesis since a higher load of *Fusobacterium* is related to disease severity [35]. From our results, patients with cancer showing a positive FIT result are more susceptible to carry *Fusobacterium*.

### Elaborating relationships between marine eukaryotes’ associations and EFs

Next, we apply kLDM on the TARA Oceans eukaryotic dataset[24] to explore associations in natural environments and compare predicted associations by kLDM with known genus-level interactions. kLDM found two clusters corresponding to two environmental conditions, Cluster 1, consisting of 168 samples with 67 OTUs, and Cluster 2, consisting of 53 samples with 26 OTUs. 41 OTUs in Cluster 2 were filtered out because the number of samples was smaller than the number of OTUs, and kLDM removed small OTUs to infer associations efficiently. EFs of the two clusters are listed in Table S7. Cluster 1 has significantly higher salinity and temperature, but lower oxygen, phosphate and silicate concentration than Cluster 2. Top 1% associations found in these two clusters, including 23 OTU-OTU and 12 EF-OTU associations in Cluster 1, and 4 OTU-OTU and 5 EF-OTU associations in Cluster 2 were analyzed, and associations with literature support are listed in **Table 2.**

**Table 2.**
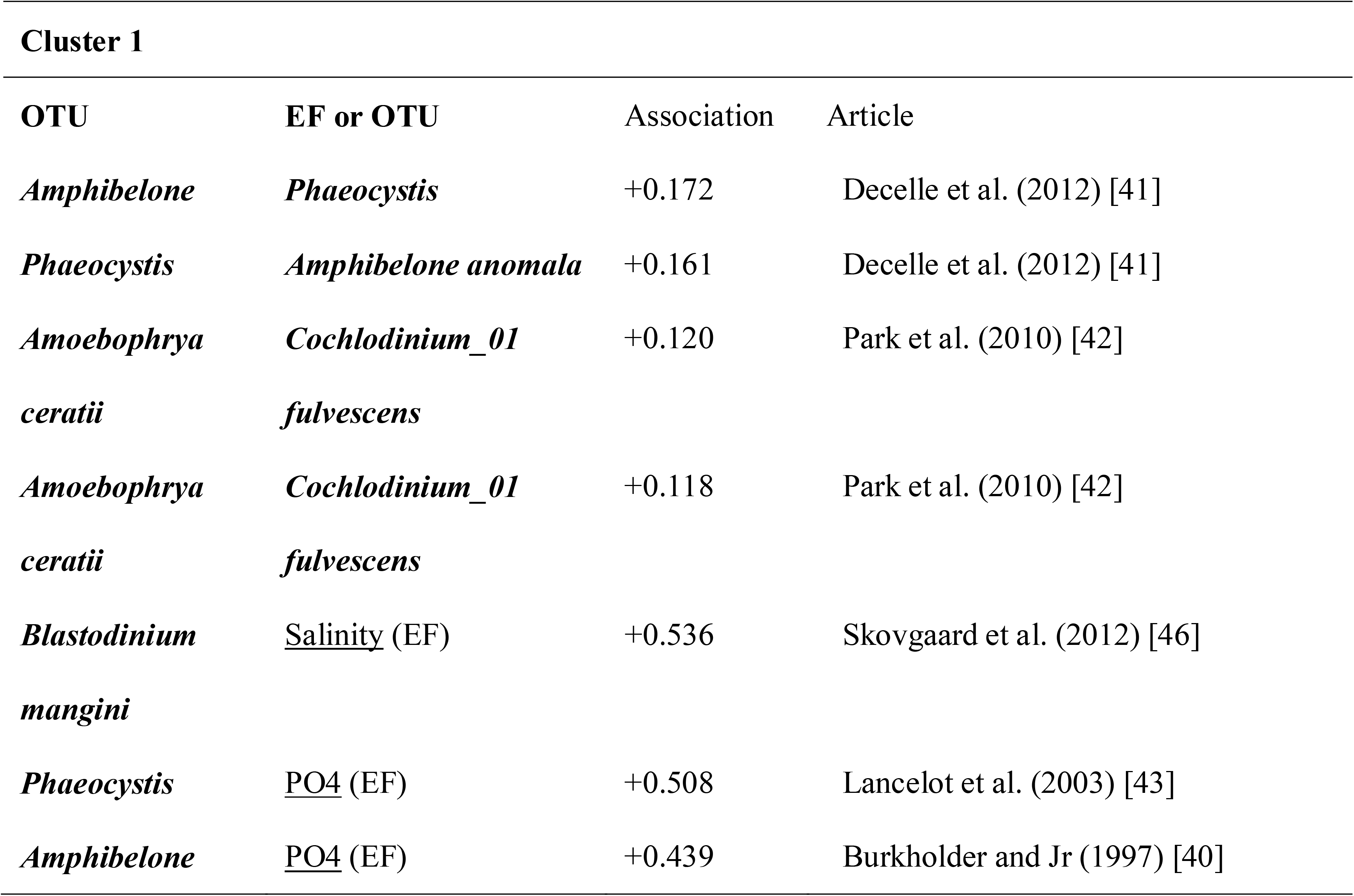
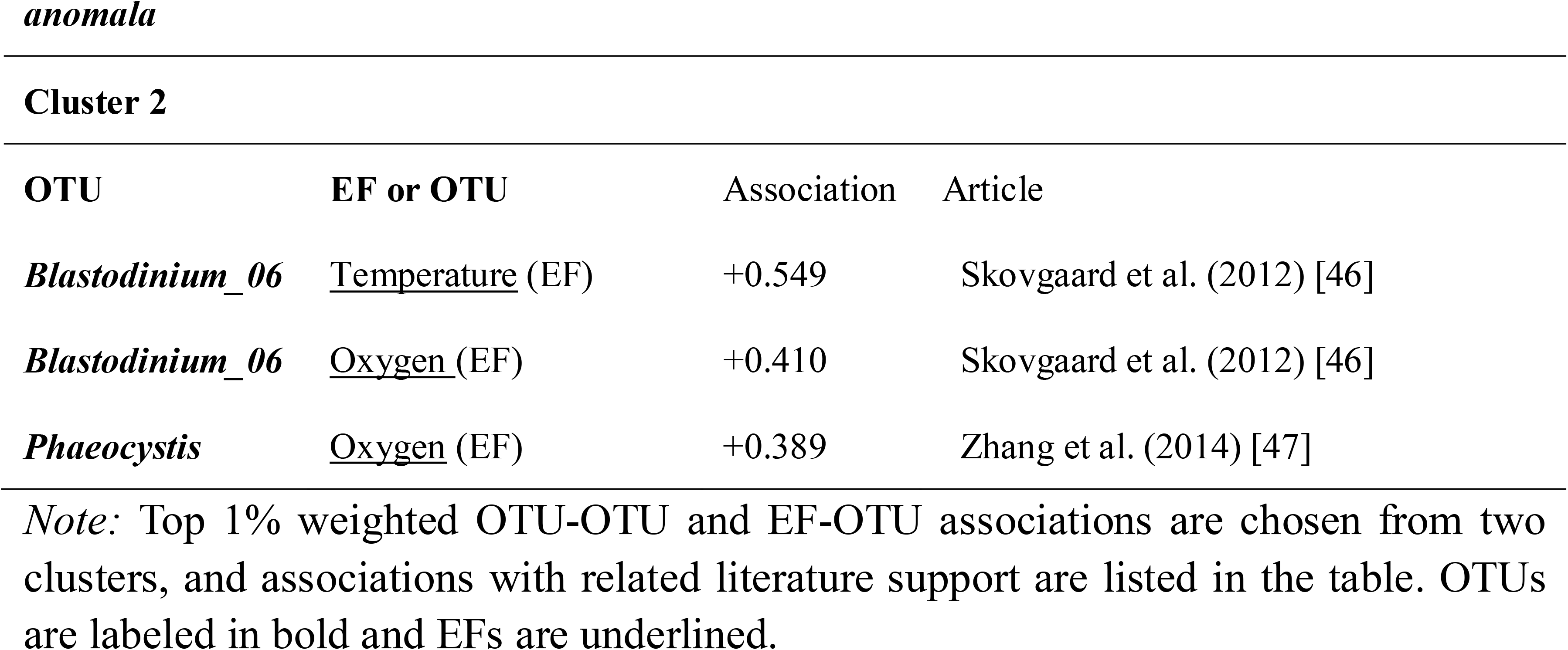
Top 1% associations of two clusters on TARA Oceans dataset with literature support.

We then matched associations with known genus-level interactions among the top-N associations within each of the two clusters discovered by kLDM. Since known interactions are at the genus-level, we regard one association between two OTUs as matched if the OTUs’ genera are identical to two corresponding genera of the known interaction. Simultaneously the results are compared with those that do not consider the environmental condition and regard all samples as one cluster, denoted as ‘Static’, by limiting kLDM to predict one set of EF-OTU and OTU-OTU association networks. From Table S8, we observe that the combined predicted associations from the two clusters are similar to that of the ‘Static’, with each cluster consisting of both common and specific OTU-OTU associations.

Different types of known associations detected by kLDM are listed in Table S9. We observe that four known associations, ‘*Phaeocystis - Amphibelone*’, ‘*Vampyrophrya - Copepoda*’, ‘*Amoebophrya - Acanthometra*’and ‘*Blastodiniaceae - Copepoda*’, may be dominant in the ocean because they are inferred from both whole dataset, and the bigger cluster (Cluster 1), and the association between *Amoebophrya* and *Protoperidiniaceae* may be strong in a specific EF condition related to Cluster 2. More specifically, the association between OTU-38 and OTU-25, belonging to the genera *Amoebophrya* and ‘Protoperidiniaceae’ respectively, only exists in Cluster 2, in which the mean values of oxygen concentration, phosphate, and silica are higher than those in Cluster 1, and the abundance levels of two OTUs are significantly higher than those in Cluster 1 (P-value < 0.05). The EF condition in Cluster 2 could be more suitable for growth of the genus *Protoperidiniaceae*, and then *Amoebophrya* would benefit from parasitism with the former. Based on results from the TARA Oceans dataset, we confirmed the effectiveness of kLDM to elaborate the relationship between OTU-OTU associations and values of EFs.

### Characterizing changes in human gut microbes’ association networks with different lifestyle factors

kLDM has the advantage of analyzing complex datasets with large samples to infer multiple EF conditions and the corresponding association networks, and we verified this capability on the American Gut project dataset[14]. We applied kLDM to cluster samples according to lifestyle-related factors and association networks, and obtained two big clusters, ‘C1’ with 6,831 samples and ‘C2’ with 5,003 samples, and one small cluster ‘C3’ with 112 samples.

Compared with ‘C1’ and ‘C2’, ‘C3’ exhibits different lifestyle patterns and microbiota distributions (Table S10, S11, and S12). ‘C1’ and ‘C2’ together contain almost all disease and healthy people, while ‘C3’ is mainly composed of IBD patients (94.64%), who make up about a quarter of all IBD patients (26.77%). Individuals in ‘C3’ have significantly higher alcohol frequency, high fat red meat frequency, and red meat frequency than those in clusters ‘C1’ and ‘C2’, but they also eat more vegetables and take vitamin D and B supplements and probiotics more frequently. However, their exercise frequency, milk substitute frequency, and milk cheese frequency are distinctly lower (Table S10). For genus-level microbial abundance, genera ‘*Prevotella*’, ‘*Ruminococcus*’, and ‘*Sutterella*’ are more abundant in ‘C3’, but ‘*Bifidobacterium*’ and ‘*Bacteroides*’ are less abundant (Table S13). If only considering IBD samples in each cluster, we observe that the IBD samples in ‘C1’ and ‘C2’ are substantially different from those in ‘C3’ based on the aforementioned diet frequencies and genus abundance (Table S14). Comparing IBD samples in ‘C1’ and ‘C2’, we observe similar lifestyle quantification values and genus abundance levels except vegetable, smoking, fruit, and home-cooked meal frequencies and the abundance of the genus ‘*Lachnospira*’.

Then the top 1% EF-OTU associations predicted by kLDM were analyzed (Table S12), and various associations among three clusters matched with published literature are shown. The positive correlation between probiotic frequency and ‘*Bifidobacteria*’ is observed in both ‘C1’ and ‘C2’, which was previously reported by Rajkumar et al. [36]. In high animal-protein diet, we observed positive correlations between red meat frequency and ‘*Bacteroides’*, and between poultry frequency and ‘*Ruminococcus*’ in ‘C1’, and between seafood frequency and ‘*Clostridiales*’ in ‘C3’, all consistent with other research [37]. Associations between high-fat red meat frequency and ‘*Bacteroides*’ in ‘C1’ and ‘*Clostridiales*’ in ‘C2’ are matched with rat study showing that intake of a high-fat diet results in disproportionately increases in propionate- and acetate-producing species such as ‘*Clostridiales*’ and ‘*Bacteroides*’[38].

## Discussion

Considering the dynamic nature of microbial interactions, we proposed a new hierarchical Bayesian model, kLDM, to infer associations among microbes and between environmental factors and microbes under different environmental conditions. We then developed two algorithms, a theoretical EM algorithm, and a more practical and efficient split-merge algorithm, to estimate both the number of environmental conditions and associations for each environmental condition simultaneously. The effectiveness of kLDM was verified on simulated datasets, as well as real TARA Oceans, CRC, and American Gut project datasets. Although we implemented kLDM with OpenMP, it can also be accelerated in MPI.

For the synthetic experiment, when we tested kLDM’s scalability (Figure S1A and S1B), AUC scores of CCLasso were the second-best in most cases, but CCLasso predicted more associations than kLDM, and its ROC curves increased slowly at the beginning and climbed higher with increase of the false positive rate. However, we think the initial high true positive rate is more crucial for scientists wanting to select candidate interactions for validation; thus kLDM has its advantage here. In addition, influence of the similarity level of EF values between different EF conditions on the performance of kLDM was further explored in detail. We plotted the relationship between AUC scores and the absolute distance (see the formula in Table S3) between EF values of two clusters (Figure S5) based on the ‘**baseline**’ dataset used in Table S1. When the distance is small (distance < 2), we observed that AUC scores of OTU-OTU and EF-OTU associations would increase rapidly with the distance, and when distance is larger than two, the power of kLDM tends to be stable. To maintain all AUC scores >0.7, the distance of EF values of two clusters should be larger than 1.0. We have also computed mean values of the inverse Simpson index *n*_*eff*_ of the synthetic datasets, and the results are listed in Table S16. Values of *n*_*eff*_ of two synthetic datasets with P=100 and 200 are ≥13 and have high compositionality [39]. Combining Table S16 with AUC scores of kLDM in Figure S1, we can see kLDM can handle high compositional effect.

On the CRC dataset, the diagnostic state of the donors is included as one EF by kLDM. The diagnostic state was classified according to colonoscopy examination and review of biopsies, instead of the microbial compositions and associations. Therefore, patients with the same diagnostic state may not have the same underlying microbial associations. In addition, cancer may have different subtypes, and each subtype may have a different association network. A patient who will potentially develop to cancer may also have a different association network from those who will not. We also test kLDM on the CRC dataset using three environmental factors excluding the diagnostic state. We found that the two clusters stayed the same, and the compositions of the diagnostic state in the two clusters didn’t change. The reason may be that the underlying microbial networks are distinct between these two clusters. In addition, we also compare kLDM with a probabilistic clustering model, called MicrobeDMM[40], which clusters sequencing samples according to the microbial composition only. From Table S17, we observe that MicrobeDMM doesn’t distinguish the healthy from the CRC samples well, and the gut samples of these two groups are mixed evenly into two clusters. In comparison, in the results of kLDM, most of the ‘Normal’ samples are clustered into one group, while the ‘Cancer’ samples to the other, which is consistent with our prior knowledge. This shows that using only microbial composition may not be sufficient, and this result, together with the result of kLDM on the synthetic dataset with partially observable EFs 1(Figure S2), proves the importance of collecting of all EFs.

On the TARA Oceans dataset, top associations found by kLDM were associated with previous studies (Table 2). For Cluster 1, the four listed OTU-OTU associations belong to two kinds of known microbial interactions. More specifically, the symbiosis between *Phaeocystis* and *Amphibelone anomala*, and the parasitism between *Amoebophrya ceratii* and *Cochlodinium_01 fulvescens* were captured by kLDM [41,42]. Associations related to values of EFs were also found. *Blastodinium mangini* tended to live in seawater with high salinity, and the concentration of PO4 affected the growth of *Phaeocystis* [43]. In addition, *Amphibelone anomala* is associated with PO4 because it has a close phylogenetic relationship with *Pfiesteria piscicida* [44], which is regulated by phosphate [45]. For Cluster 2, the parasite *Blastodinium_06* is linked to temperature and oxygen, and Skovgaard et al. [46] reported that some *Blastodinium spp.* living in warm temperatures can perform photosynthesis. The concentration of oxygen is associated with some *Phaeocystis*, and it was reported that the bloom of *Phaeocystis globose* caused oxygen depletion [47]. Because most interactions among marine microorganisms are unknown, our explanation is limited.

For the American Gut project dataset, kLDM analyzed the relationship between lifestyle factors and microbiota, as well as associations within the microbial community. The resultant three clusters show differential frequencies of lifestyle factors, the composition of microbes and OTU-OTU, and EF-OTU associations. Diet preference is among the most influential environmental factors on gut microbiome, and it can even determine microbial composition in the mammalian evolution process [48]. These are reflected again in our results. IBD patients in three clusters were analyzed, and the results indicated gut microbial heterogeneity of IBD patients. Potentially, subgroups could be classified by individuals’ lifestyle and microbial community. The concept of enterotype in gut microbial community has been discussed widely [49,50], and kLDM may be a useful tool to discover special subgroups of IBD patients.

We were also interested in studying whether IBD patients in ‘C3’ differed from other IBD patients in ‘C1’ and ‘C2’ in pathogenesis, clinical characteristics, treatment, and prognosis. Notably, although patients in ‘C3’ consume more probiotics, they still show low abundance of ‘Bifidobacterium’. Probiotic diets can induce the anti-inflammatory factor, IL-10, which improve gut microenvironment and reduce IBD symptoms [51,52]. Individuals in ‘C1’ and ‘C2’ showed low probiotics diet frequency, which agrees with studies in the literature, but IBD patients from ‘C3’ show high probiotics diet frequency (Table S8). Philpott and Girardin [53] reported that IBD patients carrying NOD2 mutations decreased transcription of IL-10. Whether patients in ‘C3’ are more likely to carry NOD2 mutations to cause the disease is suspected.

By further comparing the IBD diagnosis among three clusters in Table S15, all IBD patients in ‘C3’ are those with colonic Crohn’s disease, but such information of IBD samples in the other two clusters is missing. Patients with Crohn’s disease (CD) may lack vitamin B and D [54]; therefore, the use of vitamin B and D supplements in ‘C3’ indicates the treatment for such patients. Recently, the ‘Anti-Inflammatory’ diet, which avoids high-fat meats, has been reported to reduce symptoms in patients with IBD [55]. However, high fat red meat frequency is observed for colonic Crohn’s disease patients in ‘C3’, and adjustment in this category may help them in the future. Whether the lifestyle of patients with colonic Crohn’s disease is affected by a doctor’s advice is not included in the dataset, which limits further interpretation.

Taking into account healthy cohorts in ‘C1’ and ‘C2’, we observed that genus abundance of the IBD samples in ‘C1’ or ‘C2’ shared more similarity with that of the healthy samples when compared to the IBD samples in ‘C3’ (Table S18). Recently many microbiota-assisted models have been proposed to detect gut lesion but with mixed results [50,56,57]. Adding microbial abundance improves sensitivity but with the sacrifice of specificity. Our results indicate that such decreased specificity may result from the close microbial composition between some patients and that of healthy people. Therefore when we design microbiota-based prediction models, heterogeneity within patient samples should be considered.

While we demonstrated that kLDM is an excellent tool for biologists to interpret microbial associations under multiple environmental conditions, there are implementation challenges, as well as several possible improvements. For example, the mathematical score (e.g., EBIC score) may not be sensitive enough to separate two environmental conditions. Studies about differential gene co-expression networks may be useful for kLDM. For example, characteristics of nodes in networks, such as degree, clustering coefficient, and other mathematical measures summarizing the change in associations, may be included to build a more suitable approach to merging sub-clusters [58]. In addition, the assumption of the Gaussian distributions for the EFs may not be very suitable in the case of categorical data types, and other probabilistic distributions can be considered to model categorical meta data. The scalability of kLDM needs to be expanded further so that a large number of rare OTUs with low-occurrence can be included in the model. Prior knowledge of microbes and their interactions from known studies may also be very helpful for kLDM to reduce the complexity of association inference and improve model’s efficiency further [59]. The sample sizes of the real TARA Oceans and CRC datasets are still relatively small, which is a common phenomenon in current metagenomic studies, but we believe that more large scale datasets such as the American Gut Project and Human Microbiome Project, will be collected in the future. For these large-scale datasets, kLDM could be the perfect tool to analyze complex associations.

## Conclusions

We have proposed kLDM to infer EF conditions existing in the microbial community and predict direct EF-OTU and conditionally dependent OTU-OTU associations under every EF condition, considering compositional bias simultaneously. Compared with traditional methods that estimate static association networks, kLDM has the advantage of illuminating the influence of EFs on associations in the microbial community. The EF condition estimated by kLDM is the result of taking EFs, OTUs’ abundance, and associations in the community into account comprehensively, which can provide biologists with more insight into the heterogeneity of the microbial community and identify microbes and interactions regulated by EFs, such as nutrients, lifestyle, and health status. kLDM’s superiority is validated on both synthetic data and real datasets related to human gut and marine ecosystem. With the deepening of research on the relationship between microorganisms and human diseases, it is expected that kLDM will enable new discoveries of variation of microbes and OTU-OTU and EF-OTU associations with human health and diet.

## Authors’ contributions

YY, XW, NC and TC designed the method. YY and XW developed the algorithm and performed data analysis. YY, XW, KX, CZ, NC and TC drafted the manuscript. All authors read and approved the final manuscript.

## Acknowledgments

This work is supported by the National Natural Science Foundation of China (Nos: 61872218, 61673241, 61721003), and Tsinghua-Fuzhou Institute Research Program, Beijing National Research Center for Information Science and Technology (BNRist).

## Supplementary material

**Figure S1 AUC scores of kLDM with increasing numbers of microbes and EF conditions**

The ‘K’ is the number of environmental conditions or clusters. **A.** AUC scores with 100 microbes, 8 environmental factors and 2 clusters. Size of samples of clusters belongs to [400, 800]. **B.** AUC scores with 200 microbes, 10 environmental factors and 2 clusters. Every cluster’s number of samples range from 800 to 1600. **C.** AUC scores with 50 microbes, 5 environmental factors and 3 clusters. Number of each cluster’s samples ranges from 200 to 400. **D.** AUC scores with 50 microbes, 5 environmental factors and 4 clusters.

**Figure S2 AUC scores of kLDM when EFs with different ratios are utilized**

The ‘baseline’ dataset in **Table S3** is used and we compared the results when only 20%, 40%, 60% and 80% EFs are found in the dataset. EFs are selected orderly and set to zero because they contribute EF-OTU associations equally in the simulation experiment. Change of AUCs of OTU-OTU and EF-OTU associations of two clusters are shown.

**Figure S3 CRC-associated microbial relative abundance on two clusters**

**Figure S4 OTU-OTU association networks for cancer (Cluster 1) and healthy (Cluster 2) clusters**

Edges represent associations with absolute weight among top 1%. Red and green edges represent positive and negative associations, respectively. The width of an edge is proportional to the association’s absolute weight. Only microbes with associations in either Cluster 1 or Cluster 2 are shown. Positions of OTUs in the A and B are exactly identical and sizes of nodes are correlated with their average abundance in all samples. 5 CRC-associated microbes (Peptostreptococcus, Parvimonas, Fusobacterium, Porphyromonas and Prevotella), are arranged closely and colored with blue-green, and their associated microbes are labeled with red.

**Figure S5 The relationship between the effectiveness of kLDM and the similarity of EFs of EF conditions**

The ‘baseline’ dataset in **Table S3** is used again and we changed the distance between two clusters’ mean values of EFs from 0.1 to 10.0. For every distance, AUC scores of estimated OTU-OTU associations (‘Cluster 1 OTU-OTU’ and ‘Cluster 2 OTU-OTU’) and EF-OTU associations (‘Cluster 1 EF-OTU’ and ‘Cluster 2 EF-OTU’) of two clusters are calculated.

**Table S1 Comparison of running time and memory usage by kLDM and mLDM on single association network inference**

**Table S2 Mean values of sensitivities and specificities of top N associations in five different methods**

**Table S3 AUC scores of kLDM considering the influence of similarity of environmental conditions, EF-OTU associations and OTU-OTU associations respectively**

**Table S4 AUC scores of kLDM when EFs with different ratios are utilized and other methods with full dataset**

**Table S5 Mean values of meta data of two clusters on colorectal cancer data**

**Table S6 Composition of the diagnostic state in two clusters on colorectal cancer data**

**Table S7 Mean values of EFs of two clusters in TARA Oceans dataset**

**Table S8 Matched genus-level interactions on TARA Oceans dataset of kLDM and the ‘Static’**

**Table S9 Types of matched genus-level interactions in results of kLDM and the ‘Static’**

**Table S10 20 EFs of three clusters predicted by kLDM on American Gut project dataset**

**Table S11 Health status of people in three clusters on American Gut project dataset**

**Table S12 OTU abundance, EF-OTU and OTU-OTU associations of three clusters on the American Gut project dataset**

(1) 216 OTUs’ relative abundance of three clusters. (2) Top 1% OTU-OTU associations of ‘C1’. (3) Top 1% OTU-OTU associations of ‘C2’. (4) Top 1% OTU-OTU associations of ‘C3’. (5) Top 1% EF-OTU associations of ‘C1’. (6) Top 1% EF-OTU associations of ‘C2’ (7) Top 1% EF-OTU associations of ‘C3’.

**Table S13 Genus-level relative abundance of three clusters on the American Gut project dataset**

44 genuses’ relative abundance is calculated and P-value of the abundance difference of each genus between any two clusters is computed by the Wilcoxon rank-sum test.

**Table S14 EF values and genus-level relative abundance of IBD samples in three clusters**

(1) EF values of IBD samples in ‘C1’, ‘C2’ and ‘C3’ and the P-value of each EF between any two clusters is calculated by the t-test. (2) Genus-level relative abundance of IBD samples and the abundance difference is done by the Wilcoxon rank-sum test.

**Table S15 IBD diagnosis of IBD patients in three clusters on American Gut project dataset**

**Table S16. Values of inverse Simpson index of synthetic datasets corresponding to Figure 2 and Figure S1.**

**Table S17 Composition of the diagnostic state in two clusters estimated by MicrobeDMM on colorectal cancer data**

**Table S18 EF values and genus-level abundance of healthy and IBD samples in ‘C1’ and ‘C2’**

(1) EF values of healthy and IBD samples in ‘C1’ and ‘C2’ and the P-value is computed by t-test. (2) Genus-level relative abundance of healthy and IBD samples in ‘C1’ and ‘C2’.

