## Supplementary figures and images for "Inferring Multiple Metagenomic Association Networks based on Variation of Environmental Factors"

### Figure S1

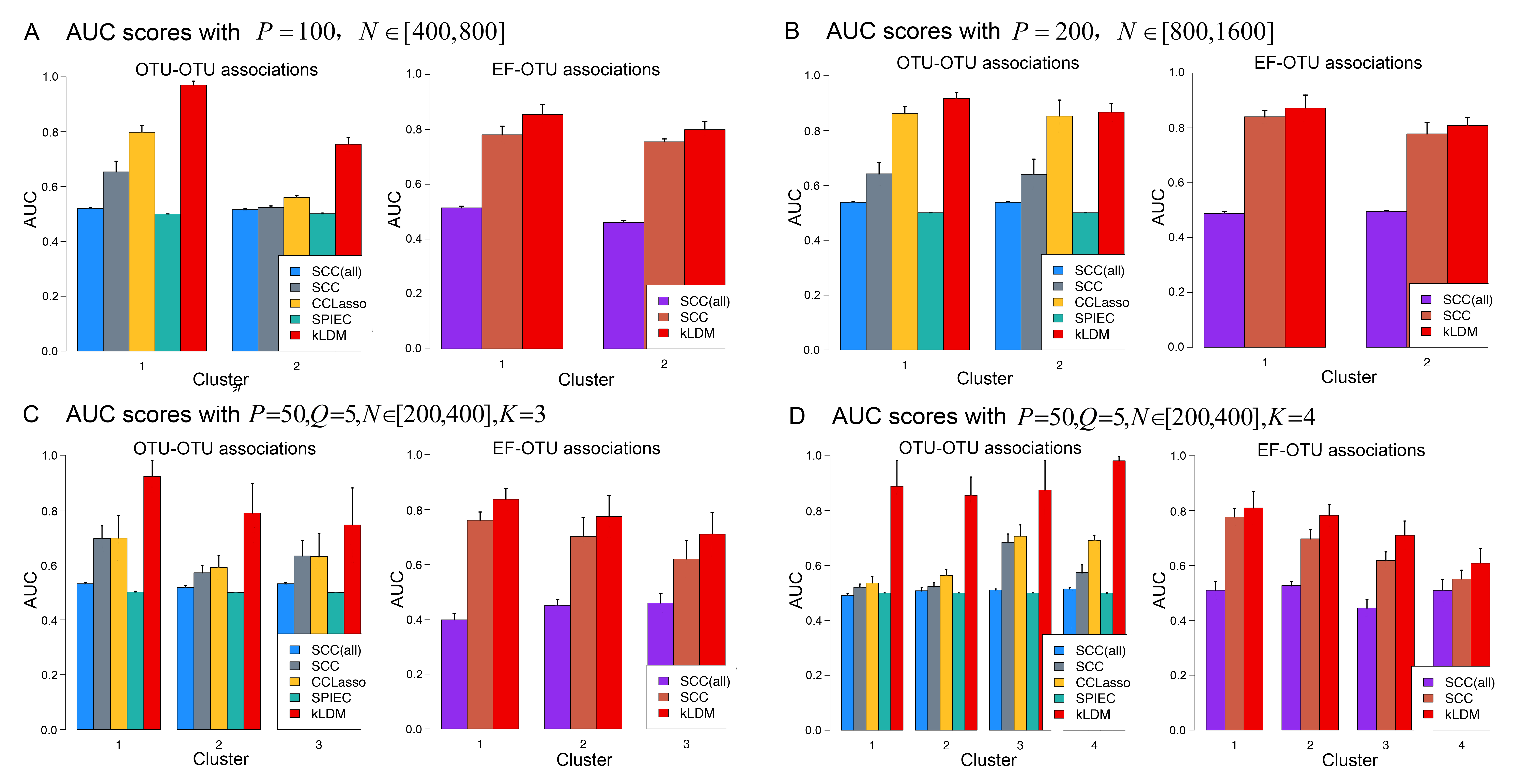

### Figure S2

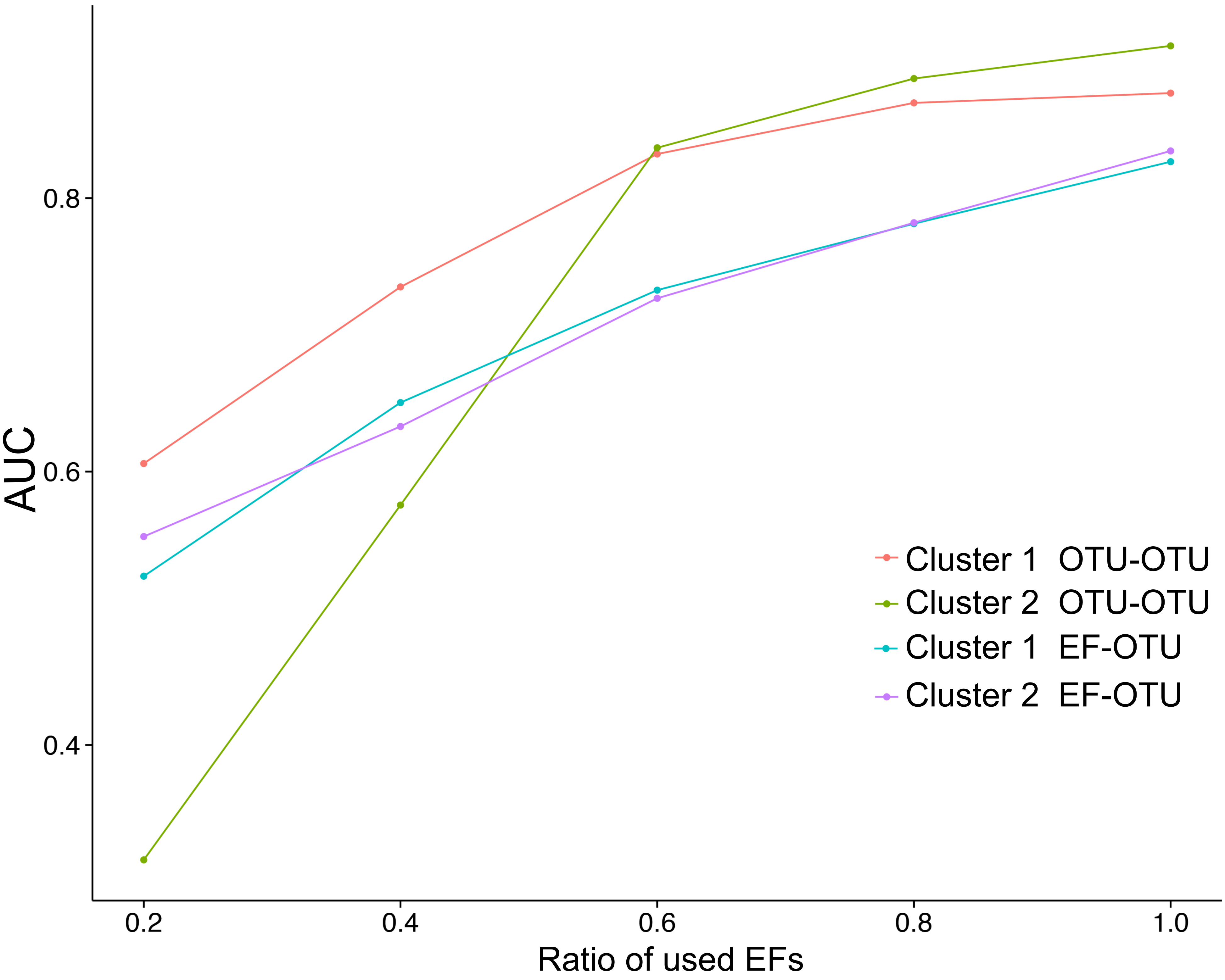

### Figure S3

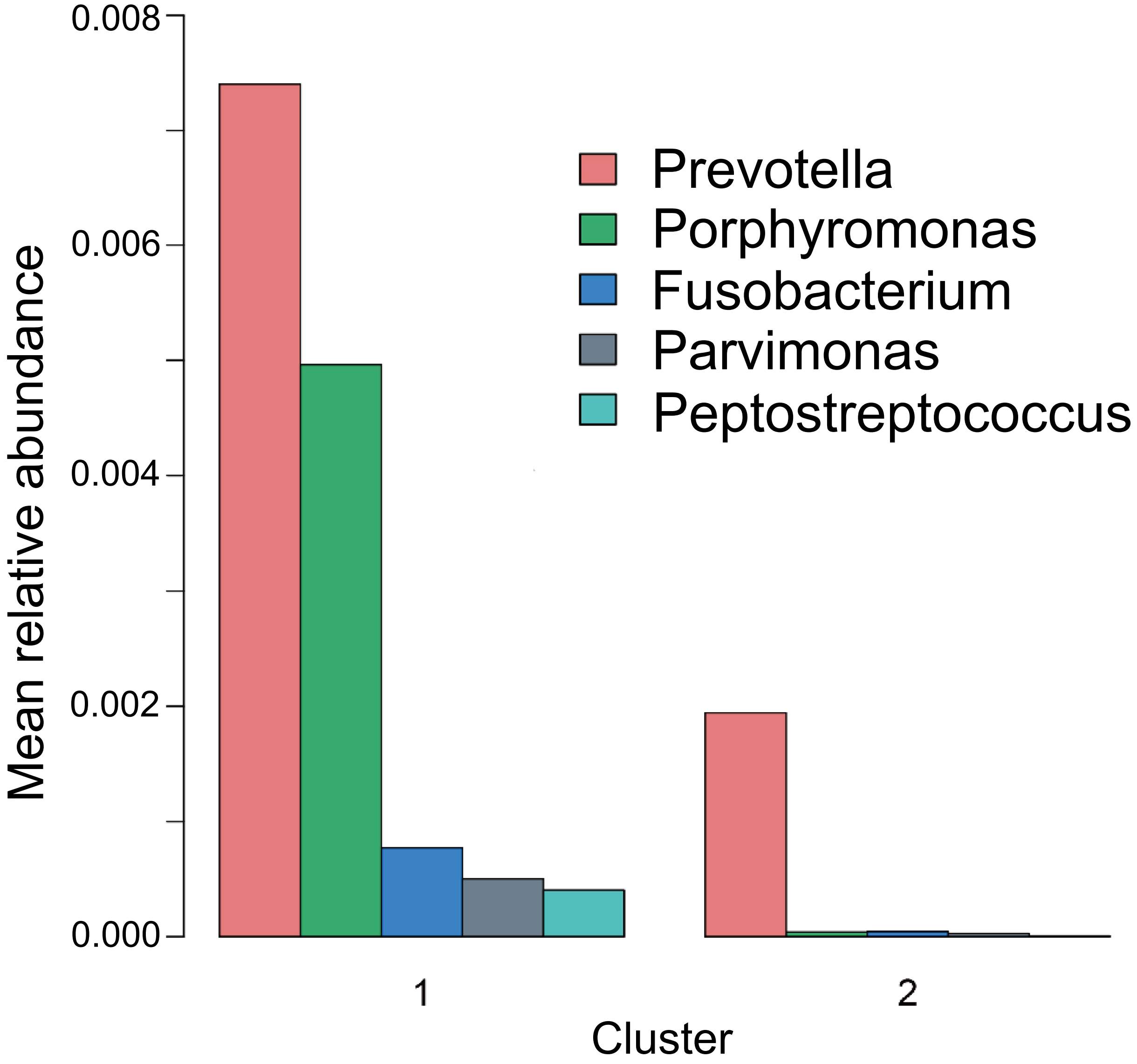

### Figure S4

A OTU-OTU associations of Cluster 1

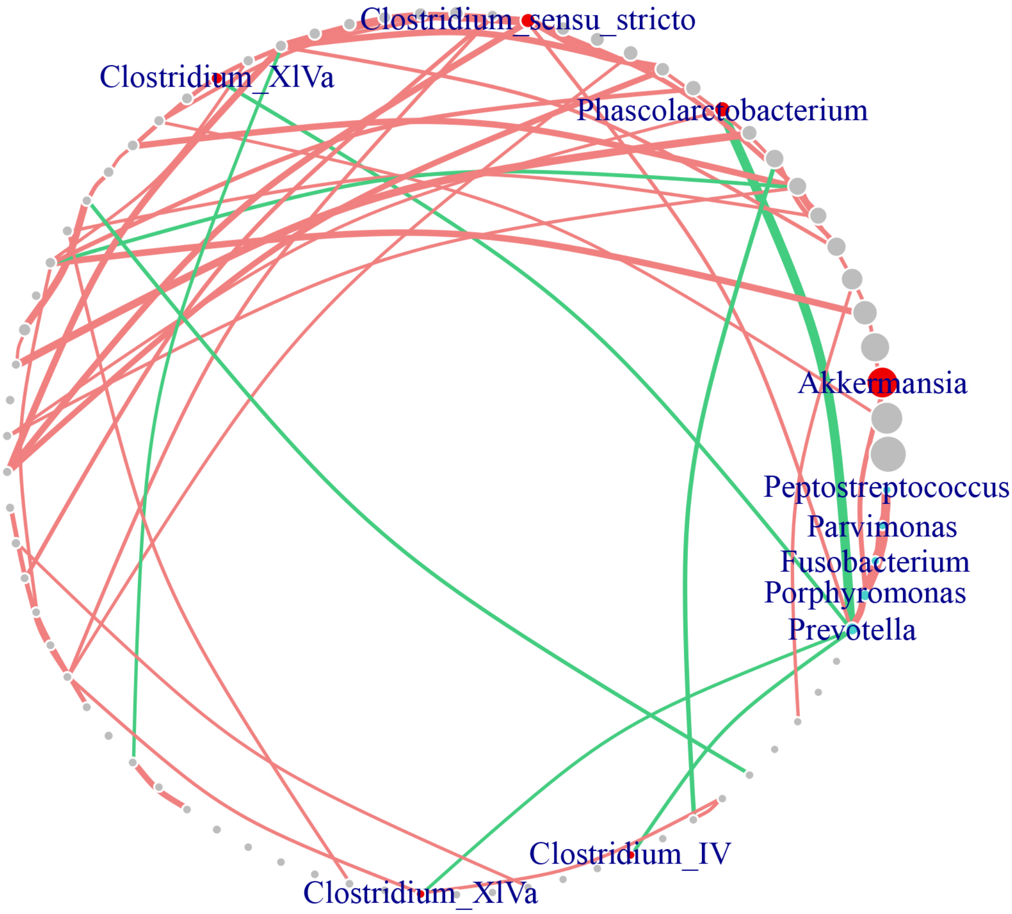


B OTU-OTU associations of Cluster 2

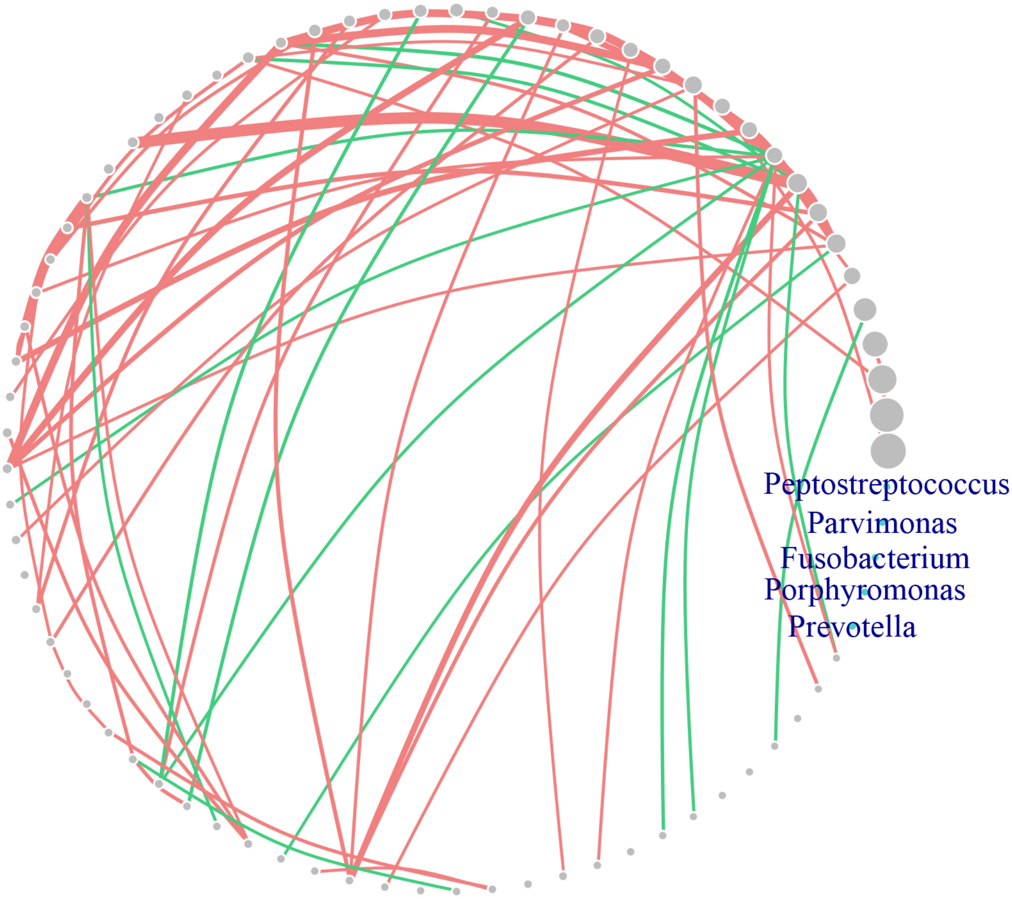

### Figure S5

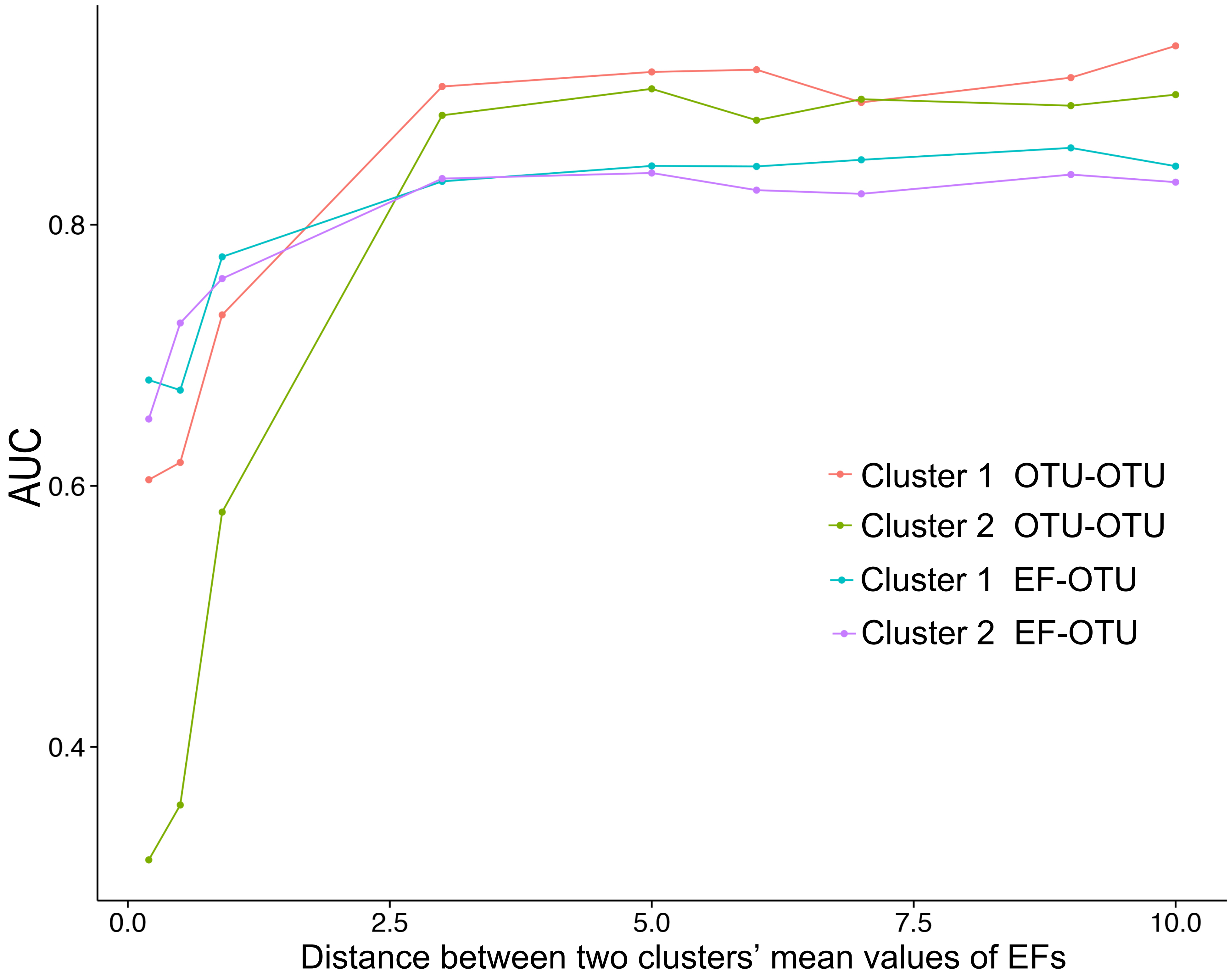
