## Supplementary material for "Inferring Multiple Metagenomic Association Networks based on Variation of Environmental Factors": Table S17

### Table S17 Composition of the diagnostic state in two clusters estimated by MicrobeDMM on colorectal cancer data

| Name | Normal | High Risk Normal | Adenoma | Advanced Adenoma | Cancer |
| --- | --- | --- | --- | --- | --- |
| Cluster 1 | 50 | 10 | 43 | 17 | 54 |
| Cluster 2 | 72 | 40 | 66 | 72 | 66 |

*Note:* The number of samples with corresponding diagnostic state in two clusters is listed.
