## Supplementary material for "Inferring Multiple Metagenomic Association Networks based on Variation of Environmental Factors": Table S16

**Table S16 Values of inverse Simpson index of synthetic datasets corresponding to Figure 2 and Figure S1**

| Synthetic datasets | Inverse Simpson index $n_{eff}$ |
| --- | --- |
| K=2, P=50, Q=5 and N ∈ [100,200] | 9.52 |
| K=2, P=50, Q=5 and N ∈ [200,400] | 9.48 |
| K=2, P=100, Q=8 and N ∈ [400,800] | 16.82 |
| K=2, P=200, Q=10 and N ∈ [800,1600] | 33.22 |

*Note:* The inverse Simpson index can be calculated by $n_{eff}=e^{-\sum_{j=1}^{P} x_{j}logx_{j}}$ where $x_{j}$ is the relative abundance of the $j^{th}$ OTU.
