## Supplementary material for "Inferring Multiple Metagenomic Association Networks based on Variation of Environmental Factors": Table S15

### Table S15 IBD diagnosis of IBD patients in three clusters on the American Gut project dataset

| Disease Name | IBD in C1 | IBD in C2 | IBD In C3 |
| --- | --- | --- | --- |
| Colonic Crohn's Disease | 7 | 1 | **106** |
| Ileal and Colonic Crohn's Disease | 5 | 1 | 0 |
| Ileal Crohn's Disease | 11 | 6 | 0 |
| Microcolitis | 3 | 0 | 0 |
| Ulcerative Colitis | 35 | 18 | 0 |
| Unknown | **163** | **40** | 0 |
| SUM | 224 | 66 | 106 |

*Note:* 'IBD' is the abbreviation for inflammatory bowel disease.
