## Supplementary material for "Inferring Multiple Metagenomic Association Networks based on Variation of Environmental Factors": Table S11

### Table S11 Health status of people in three clusters on American Gut project dataset

| Health Status | C1 (N=6831) | C2 (N=5003) | C3 (N=112) |
| --- | --- | --- | --- |
| Healthy | 55.31% / 52.68% | 67.72% / 47.24% | 5.36% / 0.08% |
| Cardiovascular Disease | 3.41% / 78.98% | 1.24% / 21.02% | 0.00% / 0.00% |
| Small Intestinal Bacterial Overgrowth | 3.62% / 73.95% | 1.74% / 26.05% | 0.00% / 0.00% |
| Mental Illness | 9.27% / 77.96% | 3.58% / 22.04% | 0.00% / 0.00% |
| Lactose Intolerant | 14.80% / 54.12% | 17.13% / 45.88% | 0.00% / 0.00% |
| Diabetes | 2.12% / 71.43% | 1.16% / 28.57% | 0.00% / 0.00% |
| Inflammatory Bowel Disease | 3.28% / 56.57% | 1.32% / 16.67% | 94.64% / 26.77% |
| Irritable Bowel Syndrome | 12.91% / 76.70% | 5.36% / 23.30% | 0.00% / 0.00% |
| Clostridium Difficile Infection | 2.56% / 84.13% | 0.66% / 15.87% | 0.00% / 0.00% |
| Cancer | 5.46% / 86.34% | 1.18% / 13.66% | 0.00% / 0.00% |
| Obesity | 9.19% / 56.94% | 9.49% / 43.06% | 0.00% / 0.00% |

*Note:* Two kinds of ratios of every health status in the cluster is shown with the format 'Ratio1 / Ratio2'. The 'Ratio1' means the ratio of number of individuals with the status in current cluster and the 'Ratio2' is the ratio in individuals with the status from whole dataset.
