## Supplementary material for "Inferring Multiple Metagenomic Association Networks based on Variation of Environmental Factors": Table S10

### Table S10 22 EFs of three clusters predicted by kLDM on American Gut project dataset

| Meta Name | C1 (N=6831) | C2 (N=5003) | C3 (N=112) | P-value  C1 vs C2 | P-value  C1 vs C3 | P-value  C2 vs C3 |
| --- | --- | --- | --- | --- | --- | --- |
| Alcohol Frequency | -0.011 (0.96) | -0.024 (1) | 1.7 (0.1) | 0.471 | 0 | 0 |
| Exercise Frequency | 0.066 (0.96) | -0.077 (1.1) | -0.59 (0.25) | 3.40E-14 | 2.34E-57 | 1.19E-44 |
| Fermented Plant Frequency | -0.14 (0.93) | 0.19 (1.1) | 0.34 (0.29) | 2.88E-68 | 6.77E-35 | 1.93E-06 |
| Frozen Dessert Frequency | -0.28 (0.69) | 0.36 (1.2) | 1.4 (0.54) | 1.42E-234 | 2.76E-60 | 8.00E-41 |
| Fruit Frequency | 0.26 (0.86) | -0.35 (1.1) | -0.31 (0.35) | 1.12E-233 | 6.52E-34 | 0.210 |
| High Fat Red Meat Frequency | -0.21 (0.79) | 0.26 (1.2) | 1.3 (0) | 1.16E-128 | 0 | 0 |
| Home cooked Meals Frequency | 0.44 (0.49) | -0.62 (1.2) | 0.78 (0.33) | 0 | 3.14E-19 | 1.94E-92 |
| Meat Eggs Frequency | 0.21 (0.84) | -0.3 (1.1) | 0.3 (0.18) | 5.39E-153 | 5.21E-05 | 8.05E-82 |
| Milk Cheese Frequency | 0.099 (0.98) | -0.12 (1) | -0.79 (0.28) | 4.59E-31 | 1.71E-68 | 3.31E-54 |
| Milk Substitute Frequency | -0.11 (0.99) | 0.17 (0.99) | -0.95 (0.26) | 1.06E-49 | 1.99E-70 | 2.31E-93 |
| Olive Oil | 0.24 (0.85) | -0.34 (1.1) | 0.4 (0.21) | 7.12E-209 | 1.04E-10 | 5.19E-88 |
| Poultry Frequency | -0.012 (0.89) | 0.019 (1.1) | -0.13 (0.39) | 0.115 | 0.003 | 0.0004 |
| Probiotic Frequency | -0.12 (1) | 0.13 (0.97) | 1.5 (0.48) | 1.10E-39 | 1.23E-64 | 2.82E-57 |
| Red Meat Frequency | -0.089 (0.88) | 0.098 (1.1) | 1.1 (0.11) | 2.60E-22 | 6.93E-249 | 3.09E-278 |
| Salted Snacks Frequency | -0.1 (0.86) | 0.17 (1.1) | -1.4 (0) | 5.16E-46 | 0 | 0 |
| Seafood Frequency | -0.061 (0.81) | 0.079 (1.2) | 0.22 (0.21) | 1.48E-12 | 9.20E-27 | 1.48E-07 |
| Smoking Frequency | -0.26 (0) | 0.36 (1.5) | -0.26 (0) | 1.30E-182 | - | 1.30E-182 |
| Vegetable Frequency | 0.41 (0.54) | -0.58 (1.2) | 0.86 (0) | 0 | 0 | 0 |
| Vitamin D Supplement Frequency | -0.069 (1) | 0.063 (0.94) | 1.4 (0) | 3.26E-13 | 0 | 0 |
| Vitamin B Supplement Frequency | -0.17 (0.98) | 0.2 (0.97) | 1.6 (0) | 1.14E-90 | 0 | 0 |
| Whole Grain Frequency | 0.12 (0.95) | -0.15 (1) | -0.75 (0.43) | 2.61E-47 | 1.14E-42 | 3.20E-28 |
| Whole Eggs | 0.11 (0.88) | -0.15 (1.1) | -0.14 (0.27) | 3.70E-43 | 6.28E-16 | 0.601 |

*Note*: Single factor two-tailed T-test is used to decide significance. The ‘-’ means that p-value of smoking frequency between ‘C1’ and ‘C3’ can’t be calculated because their values are constant and equal.
