## Supplementary material for "Inferring Multiple Metagenomic Association Networks based on Variation of Environmental Factors": Table S9

### Table S9 Types of matched genus-level interactions in results of kLDM and the 'Static'

| **'Static'** | | **kLDM Cluster1** | | **kLDM Cluster2** | |
| --- | --- | --- | --- | --- | --- |
| **Genus** | **OTU** | **Genus** | **OTU** | **Genus** | **OTU** |
| Phaeocystis, Amphibelone | (OTU-23, OTU-3) | Phaeocystis,  Amphibelone | (OTU-23, OTU-3),  (OTU-43, OTU-3) | Amoebophrya, Protoperidiniaceae | (OTU-32, OTU-25),  (OTU-38, OTU-25) |
| Vampyrophrya, Copepoda | (OTU-24, OTU-1) | Vampyrophrya, Copepoda | (OTU-33, OTU-2) |  |  |
| Amoebophrya,  Gonyaulacaceae Gonyaulax_02/03/04 | (OTU-38, OTU-12),  (OTU-39, OTU-4),  (OTU-39, OTU-12),  (OTU-54, OTU-20), | Amoebophrya ceratii, Gymnodiniaceae fulvescens | (OTU-38, OTU-14),  (OTU-59, OTU-14), |  |  |
| Amoebophrya,  Gonyaulacaceae Alexandrium_01 | (OTU-53, OTU-11),  (OTU-59, OTU-11) | Amoebophrya, Gymnodiniaceae Gymnodinium_06 | (OTU-45, OTU-38),  (OTU-59, OTU-45),  (OTU-62, OTU-45) |  |  |
| Amoebophrya,  Acanthometra | (OTU-41, OTU-38),  (OTU-52, OTU-20),  (OTU-52, OTU-38) | Amoebophrya, Acanthometra | (OTU-38, OTU-22),  (OTU-38, OTU-35)  (OTU-59, OTU-22)  (OTU-59, OTU-35) |  |  |
| Blastodiniaceae,  Copepoda | (OTU-47, OTU-2) | Blastodiniaceae,  Copepoda | (OTU-47, OTU-2) |  |  |
| Amoebophrya,  Peridiniaceae | (OTU-40, OTU-38) |  |  |  |  |
| Amoebophrya,  Protoperidiniaceae | (OTU-28, OTU-20)  (OTU-38, OTU-25) |  |  |  |  |

*Note:* Every association is shown with the format '(OTU-A, OTU-B)' and its corresponding genus-level interaction is listed at the previous column. It should be noted that multiple associations can belong to the same type of genus-level interaction. Top 120 OTU-OTU associations are considered for the 'Static' and the Cluster 1 of kLDM, and for the Cluster 2, top 20 associations are involved.
