## Supplemental Data 1 for "Inferring Multiple Metagenomic Association Networks based on Variation of Environmental Factors"

### Table S8 Matched genus-level interactions on TARA Oceans dataset of kLDM and the 'Static'

| Method | kLDM Cluster1 | kLDM Cluster2 | 'Static' |
| --- | --- | --- | --- |
| MG@Top 10 | 2 | 0 | 2 |
| MG@Top 20 | 2 | 2 | 4 |
| MG@Top 40 | 5 | - | 6 |
| MG@Top 60 | 7 | - | 8 |
| MG@Top 80 | 8 | - | 9 |
| MG@Top 100 | 9 | - | 13 |
| MG@Top 120 | 13 | - | 15 |

*Note:* ‘MG@Top N’ represents matched known genus-level interactions' number among top N predicted associations. the flag ‘-’ corresponds the entry where the number of predictions is < N. Results of two clusters of kLDM are listed separately.
