## Supplementary material for "Inferring Multiple Metagenomic Association Networks based on Variation of Environmental Factors": Table S7

### Table S7 Mean values of EFs of two clusters in TARA Oceans dataset

| EF name | Mean value in Cluster 1 | Mean value in Cluster 2 | P value |
| --- | --- | --- | --- |
| Depth | 39.66 | 35.71 | 0.59 |
| Salinity | 36.64 | 35.41 | 6.78E-11 |
| Temperature | 23.07 | 18.72 | 9.21E-4 |
| Oxygen | 195.48 | 233.56 | 1.47E-6 |
| PO4 | 0.36 | 0.62 | 0.02 |
| Si | 1.98 | 7.65 | 0.03 |
| Chlorophyll | 0.09 | 0.12 | 0.29 |
| Depth of chlorophyll maximum | 61.01 | 71.87 | 0.11 |
| Depth of maximum Brunt Väisälä frequency | 83.30 | 90.15 | 0.39 |
| Depth of maximum oxygen concentration | 34.72 | 160.98 | 2.67E-6 |
| Depth of minimum oxygen concentration | 297.80 | 553.48 | 1.32E-18 |
| Sunshine duration | 682.07 | 788.77 | 2.91E-6 |
| Moon phase | 0.13 | 0.03 | 0.04 |
| Maximum Lyapunov exponent | 0.026 | 0.056 | 8.05E-4 |
| Residence time | 9.11 | 14.32 | 0.09 |
| Latitude | 0.06 | -0.28 | 7.54E-13 |
| Longitude | -0.20 | -0.05 | 0.01 |

*Note:* Single factor two-tailed t-test is used to decide significance.
