## Supplementary material for "Inferring Multiple Metagenomic Association Networks based on Variation of Environmental Factors": Table S6

### Table S6 Composition of the diagnostic state in two clusters on colorectal cancer data

| Name | Normal | High Risk Normal | Adenoma | Advanced Adenoma | Cancer |
| --- | --- | --- | --- | --- | --- |
| Cluster 1 | 19 | 9 | 48 | 34 | **108** |
| Cluster 2 | **103** | 41 | 61 | 55 | 12 |

*Note:* The number of samples with corresponding diagnostic state in two clusters is listed.
