## Supplementary material for "Inferring Multiple Metagenomic Association Networks based on Variation of Environmental Factors": Table S5

### Table S5 Mean values of meta data of two clusters on colorectal cancer data

| **Meta Name** | **Cluster 1** | **Cluster 2** | **P-value** |
| --- | --- | --- | --- |
| FIT | 531.15 | 8.47e-7 | 7.18e-22 |
| Age | 61.81 | 59.05 | 0.013 |
| Gender (Female) | 46.89% | 53.31% | 0.066 |

*Note:* Mean values of FIT, age and gender are calculated and single factor two-tailed T-test is used to decide significance. 'FIT' stands for fecal immunochemical test results.
