## Supplementary material for "Inferring Multiple Metagenomic Association Networks based on Variation of Environmental Factors": Table S4

### Table S4 AUC scores of kLDM when EFs with different ratios are utilized and other methods with full dataset

| **Method** | **Cluster1 OTU-OTU** | **Cluster1 EF-OTU** | **Cluster2 OTU-OTU** | **Cluster2 EF-OTU** |
| --- | --- | --- | --- | --- |
| **SCC(all)** | 0.69$\pm$0.02 | 0.52$\pm$0.02 | 0.50$\pm$0.00 | 0.45$\pm$0.02 |
| **CCLasso** | 0.59$\pm$0.04 | - | 0.70$\pm$0.02 | - |
| **SPIEC** | 0.50$\pm$0.00 | - | 0.50$\pm$0.00 | - |
| **SCC** | 0.55$\pm$0.05 | 0.76$\pm$0.09 | 0.55$\pm$0.01 | 0.73$\pm$0.03 |
| **kLDM(100%)** | 0.88$\pm$0.11 | 0.83$\pm$ 0.08 | 0.91$\pm$0.04 | 0.83$\pm$0.03 |
| **EF (80%)** | 0.86$\pm$0.11 | 0.78$\pm$0.04 | 0.88$\pm$0.08 | 0.78$\pm$0.03 |
| **EF (60%)** | **0.83**$\pm$**0.11** | **0.73**$\pm$**0.05** | **0.84**$\pm$**0.08** | **0.72**$\pm$**0.05** |
| **EF (40%)** | 0.74$\pm$0.10 | 0.65$\pm$0.07 | 0.58$\pm$0.20 | 0.63$\pm$0.04 |
| **EF (20%)** | 0.61$\pm$0.03 | 0.52$\pm$0.04 | 0.32$\pm$0.08 | 0.55$\pm$0.03 |

*Note:* Results of kLDM matched with the **Figure S5** and four other methods’ results on the whole dataset are listed. The ‘EF (20%)’ means 20% EFs are employed by kLDM to infer association networks. The AUC score is formatted with 'mean value $\pm$ standard deviation'.
