## Supplementary material for "Inferring Multiple Metagenomic Association Networks based on Variation of Environmental Factors": Table S2

**Table S2 Mean values of sensitivities and specificities of top N associations in five different methods**

| **Top@N** | **Method** | **Cluster1 OTU-OTU** | | **Cluster1 EF-OTU** | | **Cluster2 OTU-OTU** | | **Cluster2 EF-OTU** | |
| --- | --- | --- | --- | --- | --- | --- | --- | --- | --- |
|  |  | **Sensitivity** | **Specificity** | **Sensitivity** | **Specificity** | **Sensitivity** | **Specificity** | **Sensitivity** | **Specificity** |
| 20 | SCC(all) | 0.022 | 0.987 | 0.098 | 0.924 | 0.016 | 0.982 | 0.058 | 0.916 |
|  | SCC | 0.028 | 0.989 | 0.375 | 0.985 | 0.019 | 0.997 | 0.431 | 0.874 |
|  | CCLasso | 0.044 | 0.997 | - | - | 0.019 | 0.996 | - | - |
|  | SPIEC | 0.000 | 1.000 | - | - | 0.000 | 1.000 | - | - |
|  | **kLDM** | **0.311** | **0.995** | **0.357** | **0.981** | **0.290** | **0.995** | **0.347** | **0.968** |
| 40 | SCC(all) | 0.036 | 0.969 | 0.167 | 0.841 | 0.032 | 0.964 | 0.144 | 0.837 |
|  | SCC | 0.058 | 0.979 | 0.716 | 0.654 | 0.037 | 0.990 | 0.924 | 0.085 |
|  | CCLasso | 0.083 | 0.991 | - | - | 0.036 | 0.987 | - | - |
|  | SPIEC | 0.000 | 1.000 | - | - | 0.000 | 1.000 | - | - |
|  | **kLDM** | **0.455** | **0.987** | **0.580** | **0.932** | **0.445** | **0.985** | **0.518** | **0.904** |
| 60 | SCC(all) | 0.052 | 0.953 | 0.271 | 0.766 | 0.047 | 0.944 | 0.223 | 0.757 |
|  | SCC | 0.097 | 0.974 | 1.000 | 0.000 | 0.150 | 0.888 | 1.000 | 0.000 |
|  | CCLasso | 0.114 | 0.982 | - | - | 0.054 | 0.978 | - | - |
|  | SPIEC | 0.000 | 1.000 | - | - | 0.000 | 1.000 | - | - |
|  | **kLDM** | **0.504** | **0.983** | **0.709** | **0.863** | **0.531** | **0.973** | **0.621** | **0.828** |

*Note:* This comparison was done on synthetic datasets with K = 2 clusters, P = 50 microbes, Q = 5 environmental factors, and N∈[100, 200] samples corresponding to **Figure 2**. 'Top@N' means the number of top associations estimated by five methods. The results of kLDM were shown in bold for the convenience of comparison.
