## Supplementary for "Inferring Multiple Metagenomic Association Networks based on Variation of Environmental Factors"

### Sparse association inference of kLDM in theory.

The generative model can be solved theoretically via Expectation Maximization (EM) algorithm and maximum a posteriori (MAP) estimation for the latent variable
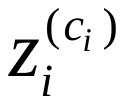
. We assume that associations among microbes and between environmental factors and microbes are sparse and can be inferred by kLDM with sparsity constraints. The logarithm of posterior distribution of the latent variables
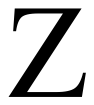
 is below,

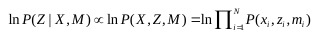

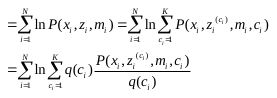

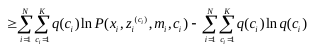

where
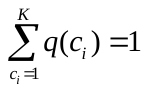
 and only when
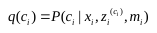
, the inequality holds according to the Jensen’s inequality. Suppose
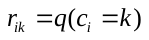
, then in the E-step, values of
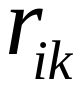
 for
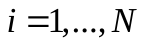
 and
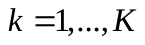
 are calculated where

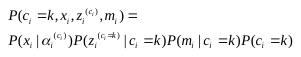

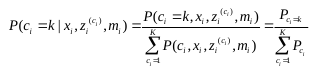

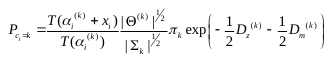

with
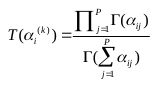
,
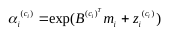
,
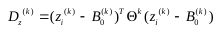
 and
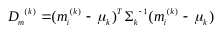
.

In the M-step, we minimize following problems:

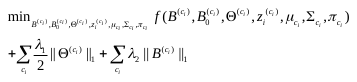

where
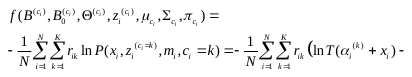

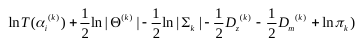
 . It should be noted that
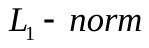
 penalties are added on both
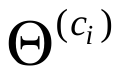
 and
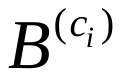
 with positive parameters
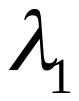
 and
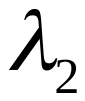
, which can help to resolve overfitting when the number of unknown parameters are much larger than the samples’ size. Then the objective function can be optimized by estimating parameters
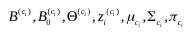
alternately.

For
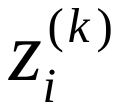
 , we minimize the objective function with respect to
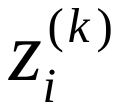
 using the L-BFGS algorithm[1] and the derivative of
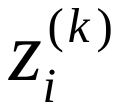
is as below,

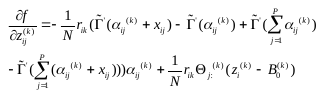

where
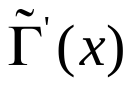
is the digamma function of the variable
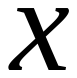
 and
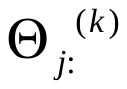
 is the
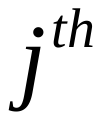
 row of the matrix
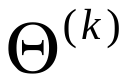
. For
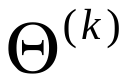
, the objective function is
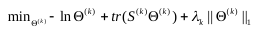
 and the inverse covariance matrix can be estimated efficiently by the QUIC method[2] with
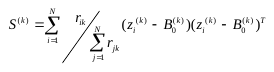
. For
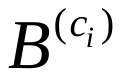
, the proximal quasi-Newton approach[3] can minimize the objective function with
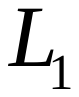
 penalty and the derivative of
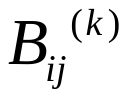
is needed as below,

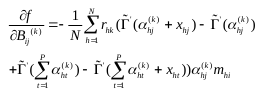

. Then
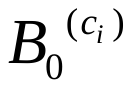
 is calculated based on
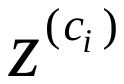
with
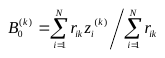
 . Other variables
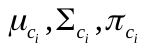
can be computed directly as
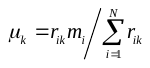
 ,
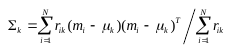
 and
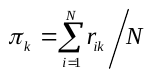
 . In order to decide the number of clusters and the best suitable values of penalty parameters, usually multiple values of
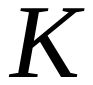
,
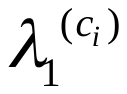
 and
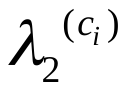
are set and the best model is selected via comparing the EBIC scores[4] among results.
