## Supplementary material for "Inferring Multiple Metagenomic Association Networks based on Variation of Environmental Factors": Table S1

### Table S1 Comparison of running time and memory usage by kLDM and mLDM on single association network inference

| Dataset  (Hub graph) | kLDM  (4 cores) | | kLDM  (8 cores) | | mLDM  (Rcpp) | | mLDM  (R language) | |
| --- | --- | --- | --- | --- | --- | --- | --- | --- |
|  | Time | Memory | Time | Memory | Time | Memory | Time | Memory |
| P=50, Q=5, N=500 | 10min  51s | 192MB | 5min  49s | 175MB | 23min  19s | 226MB | 1h 13min | 300MB |

*Note:* A synthetic dataset with 50 OTUs, 5 EFs and 500 samples was constructed and all programs were run on a server with CentOS 7.4 operating system and Intel(R) Xeon(R) E5-2680 v3 @ 2.50GHz CPUs. The ‘mLDM (R language)’ is implemented in R language, the ‘mLDM (Rcpp)’ rewrites original R functions in C++ language and integrates codes into a R package. In kLDM, we use the pure C++ codes to infer association networks and add the ability of multithread processing.
