## Supplementary material for "Inferring Multiple Metagenomic Association Networks based on Variation of Environmental Factors": Table S3

### Table S3 AUC scores of kLDM considering the influence of similarity of environmental conditions, EF-OTU associations and OTU-OTU associations respectively

| $\left\vert\boldsymbol{\mu}_{\boldsymbol{1}\boldsymbol{j}}\boldsymbol{-}\boldsymbol{\mu}_{\boldsymbol{2}\boldsymbol{j}} \right\vert\boldsymbol{=}$ | Cluster1 OTU-OTU | Cluster1 EF-OTU | Cluster2 OTU-OTU | Cluster2 EF-OTU |
| --- | --- | --- | --- | --- |
| 1.0 baseline | $0.78\pm0.07$ | $0.79\pm0.08$ | $0.69\pm0.14$ | $0.80\pm0.06$ |
| 1.0 same EF-OTU | $0.74\pm0.06$ | $0.83\pm0.04$ | $0.66\pm0.10$ | $0.78\pm0.04$ |
| 1.0 same OTU-OTU | $0.74\pm0.07$ | $0.80\pm0.08$ | $0.77\pm0.07$ | $0.78\pm0.05$ |
| 1.5 baseline | $0.89\pm0.08$ | $0.83\pm0.04$ | $0.87\pm0.07$ | $0.83\pm0.04$ |
| 1.5 same EF-OTU | $0.88\pm0.08$ | $0.85\pm0.02$ | $0.82\pm0.07$ | $0.79\pm0.04$ |
| 1.5 same OTU-OTU | $0.87\pm0.08$ | $0.85\pm0.03$ | $0.88\pm0.07$ | $0.83\pm0.03$ |
| 2.0 baseline | $0.92\pm0.04$ | $0.84\pm0.08$ | $0.86\pm0.04$ | $0.83\pm0.09$ |
| 2.0 same EF-OTU | $0.92\pm0.04$ | $0.85\pm0.03$ | $0.87\pm0.08$ | $0.82\pm0.04$ |
| 2.0 same OTU-OTU | $0.92\pm0.04$ | $0.84\pm0.03$ | $0.89\pm0.03$ | $0.84\pm0.02$ |

*Note:* Datasets with 50 microbes, 5 environmental factors and 2 clusters each with 200-400 samples are generated. Then '**baseline**' datasets are produced by changing the distance between mean values of two clusters' environmental factors. $|\mu_{1j}-\mu_{2j}|=1$ indicates that the distance of every element of mean values of two clusters' environmental factors ($\mu_{1}\mathrm{and}\mu_{2}$) is 1.0. The 'same **EF-OTU**' or 'same **OTU-OTU**' dataset is based on the '**baseline**' via merely limiting the equality of EF-OTU associations or OTU-OTU associations separately. The name on the left column, '***1.5 same EF-OTU***' means that $|\mu_{1j}-\mu_{2j}|=1.5$ and two clusters' real EF-OTU associations are identical. The AUC score is listed with the format 'mean value $\pm$ standard deviation'.
